# Expansion and collapse of VEGF diversity in major clades of the animal kingdom

**DOI:** 10.1101/2022.09.19.507521

**Authors:** Khushbu Rauniyar, Honey Bokharaie, Michael Jeltsch

## Abstract

The vascular endothelial growth factor (VEGF) family comprises in vertebrates five or six members: VEGF(-A), PlGF, VEGF-B, VEGF-C, VEGF-D, and – in venomous reptiles – VEGF-F. They fulfill mainly functions for the blood and lymphatic vascular systems. Together with the platelet-derived growth factors (PDGF-A to -D), they form the PDGF/VEGF subgroup among cystine-knot growth factors. Despite an absent vascular system in most invertebrates, PDGF/VEGF-like molecules have been found in, e.g., *Drosophila melanogaster* and *Caenorhabditis elegans*. The evolutionary relationship between PDGF and VEGF growth factors has only been addressed by older analyses, which were limited by the sparse sequencing data at the time. Here we perform a comprehensive analysis of the occurrence of PDGF/VEGF-like growth factors (PVFs) throughout all animal phyla and propose a likely phylogenetic tree. The three major vertebrate whole genome duplications play a role in the expansion of PDGF/VEGF diversity, but several limited duplications are necessary to account for the temporal pattern of emergence. The phylogenetically oldest PVFs likely featured a C-terminus with a BR3P signature, a hallmark of the modern-day lymphangiogenic growth factors VEGF-C and VEGF-D. Some of the younger *VEGF* genes appeared completely absent in some clades, e.g., functional *VEGFB* genes in the clade Archosauria, which includes crocodiles, birds, and other dinosaurs, and *pgf* in amphibians. The lack of precise counterparts for human genes poses limitations but also offers opportunities for research using organisms that diverge considerably from humans if the goal is to understand human physiology.

**Graphical abstract:** Sources for the graphical abstract:
326 MYA and older [1]
272-240 MYA [2]
235-65 MYA [3]

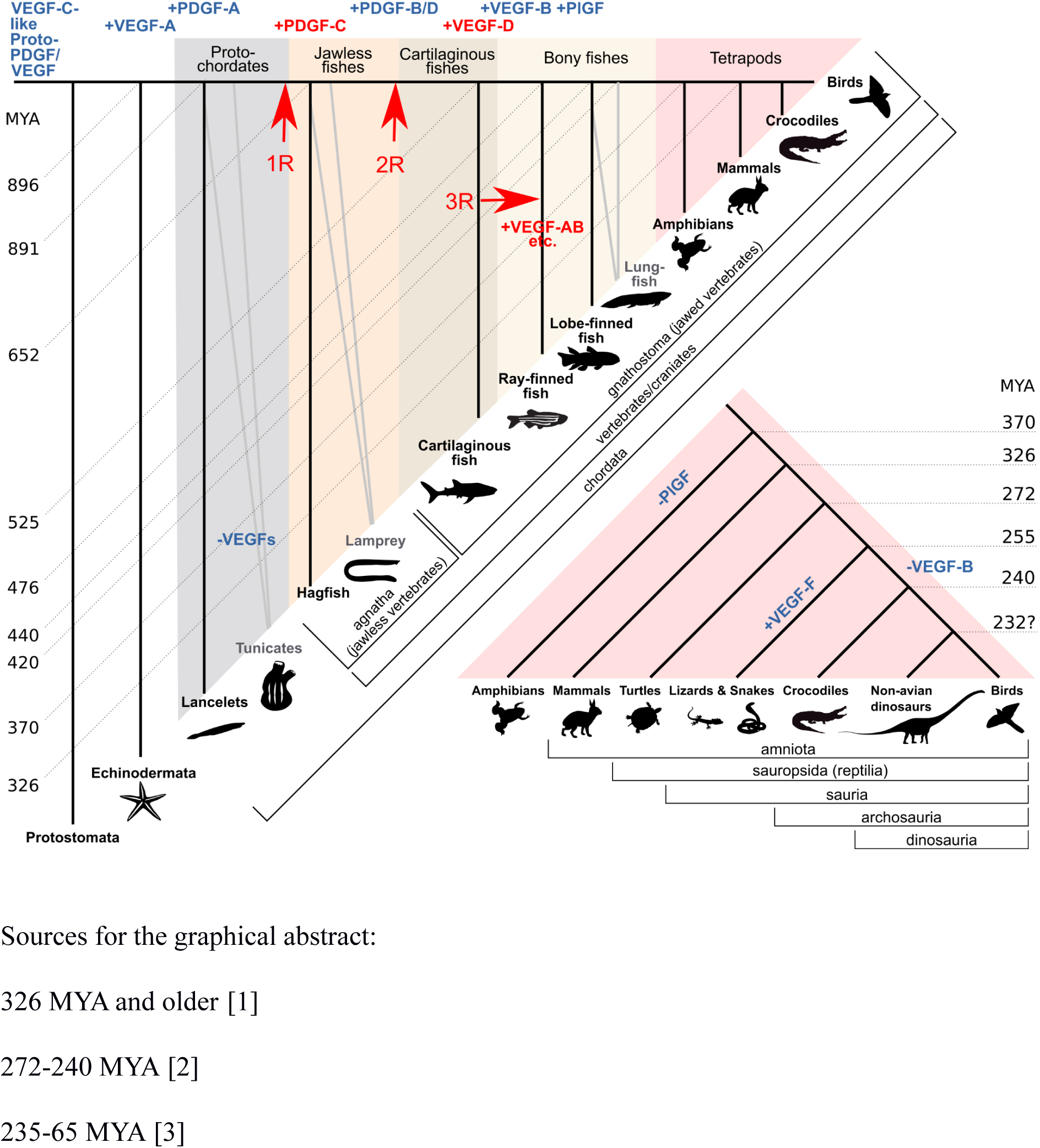

## Introduction

In biomedical research, model organisms are often used with the ultimate goal to understand the human organism. While the mouse is the most common model organism, other species can have specific advantages. Compared to the mouse, *Drosophila melanogaster* has, for example, a much shorter generation cycle, and in zebrafish or chickens, embryonic development can be directly and continuously visually observed.

Extrapolating molecular biomedical research results from a given model organism to humans is facilitated when the human proteins of interest have a corresponding counterpart (ortholog) in the model organism. Therefore, the choice of mouse as the most common model organism for biomedical research is understandable since, for many protein families, there is a 1:1 relationship of genes and proteins between *Mus musculus* and *Homo sapiens*. While there are stunning exceptions (e.g., among the kallikrein-like peptidases)[4], an early estimate found that less than 2% of mouse genes did not have a human counterpart [5]. We were interested in whether this assumption of a 1:1 relationship holds true for the vascular endothelial growth factor (VEGF) family between humans and frequently used model organisms.

### The VEGF protein family

The VEGF protein family is a highly conserved subgroup within the cystine-knot superfamily of growth factors, which share a conserved cystine knot structure, where six conserved cysteine residues are linked by three disulfide bridges such that two bridges form a ring through which the third bridge passes [6]. In vertebrates, the VEGFs are primarily involved in angiogenesis and lymphangiogenesis, the two basic mechanisms of how blood and lymphatic vessels grow [7, 8]. The VEGFs signal via the VEGF receptors (VEGFR-1, VEGFR-2, and VEGFR-3), which form a subgroup among the receptor tyrosine kinases (RTKs) which is characterized by seven extracellular Ig-like domains and an intracellular, split kinase domain [9]. Of all VEGFs, VEGF-A was discovered first and soon shown to be of paramount importance for the development of blood vessels as it was the first-ever gene for which a heterozygous deletion was found to be embryonically lethal [10, 11]. Unlike VEGF-A, even the complete loss of the subsequently discovered placenta growth factor (PlGF) and VEGF-B were reasonably well tolerated in mice [12–14]. Thus VEGF-A has been regarded as the primary, most important VEGF. VEGF-C and VEGF-D, first described in 1996, form a distinct subset within the VEGF family due to their unique structure and function [15, 16]. They feature long, distinct N- and C-terminal propeptides, require multiple proteolytic cleavages for activation, and interact with VEGFR-3, which results in their exclusive ability to directly stimulate the growth of lymphatic vessels in vivo [17–19].

### VEGF-E and VEGF-F

Two further VEGF family members have been described: VEGF-E and VEGF-F. VEGF-E is the collective name for VEGF-like molecules encoded by viruses [20], and VEGF-F denotes a group of VEGF-like molecules identified from the venom of snakes, starting with the aspic viper in 1990 [21, 22]. In the following years, many venomous snakes were shown to feature similar VEGF-like molecules [23].

VEGF-E sequences are encoded in the genomes of parapoxviruses, and their existence has been tentatively explained by a single horizontal host-to-virus gene transfer event [24], similar to how the oncogenic v-sis (a homolog of PDGF-B) is thought to have been acquired from its simian host [25].

### Invertebrate VEGFs

In invertebrates, PDGF/VEGF-like molecules have been identified, many of which are referred to as PDGF/VEGF-like growth factors (PVFs) because their exact relationship to the VEGF and PDGF growth factors appeared unclear. Together, the VEGFs and PDGFs form the PDGF/VEGF superfamily. Likely, PDGFs appeared first in the chordate lineage after the divergence from echinoderms [26, 27]. Correspondingly, their cognate receptors split before the chordates/tunicates divergence into class III and class V RTKs [28]. The fundamental biological change associated with this evolutionary period was the pressurization of the vascular system, and PDGFs are central players in the stabilization of vessels via mural and smooth muscle cells [29]. Although many PVFs can be identified from invertebrate genomic sequences, only a few have been subjected to functional analysis, including the *D. melanogaster* PVFs and *C. elegans* PVF-1. In vertebrates, VEGF receptors are expressed by cells of vascular endothelial and hematopoietic lineages, and the molecular integration of the immune and the vascular systems appears to be conserved also in invertebrates [30]. A molecular manifestation of this integration is the essential expression of the VEGF receptor-2 (VEGFR-2) by the precursor(s) of both hemopoietic and vascular endothelial lineages [31]. Unsurprisingly, the VEGFR-2 ligand VEGF-C has been shown to be important for various steps in hematopoiesis [32–34].

Opposed to this, no coherent image of the role of PVF signaling in invertebrates has emerged so far. Three separate studies involve *D. melanogaster* PVFs in immune function [35], survival of glia and neural progenitor cells [36], and mobilization of storage fat from adipocytes [37], while the *C. elegans* PVF-1 [38] functions reportedly as a repressor of Netrin signaling in the patterning of the sensillae of the male tail [39].

Five studies describe the phylogenetic relationships within the VEGF family of growth factors [40–43, 26]. However, the studies by Holmes/Zachary and Kasap suffer from a lack of comprehensive data, which was not available in 2005, while the results by Dormer and Beck and He are difficult to parse as they lack sufficient biological context. Thus, we performed a comprehensive analysis of the occurrence of PDGF- and VEGF-like sequences in the animal kingdom and proposed, based on our phylogenetic analyses, a likely evolutionary pathway, integrating it with the biological function of the PDGF/VEGF family members.

## Results

### Coverage

49992 hits were generated using 676 individual blastp searches for homologs of PDGF/VEGF family members. The searches were generated by combining 13 query sequences with 52 animal clades (see Supplementary Figure S1 for the bioinformatics workflow). 8666 of the blast hits were unique. 90.5% of these hits could be programmatically classified as members of the PDGF/VEGF protein family based on explicit manual annotation of the sequence or the PDGF motif (https://www.ncbi.nlm.nih.gov/Structure/cdd/cddsrv.cgi?uid=cd00135). The remaining 9.5% were manually examined and classified. The majority of programmatically unclassified hits appeared to be homologs of the Balbiani ring 3 protein (BR3P), to which the C-terminal domain of VEGF-C bears a striking homology [15]. A very small number of partial sequences were too short to allow classification, in which case they were excluded from further analysis. A summary of the results is shown in Figure 1, and a complete table of all hits is shown as Supplementary Table 1, and an interactive online version of the table is available at https://mjlab.fi/phylo).

**Figure 1.**
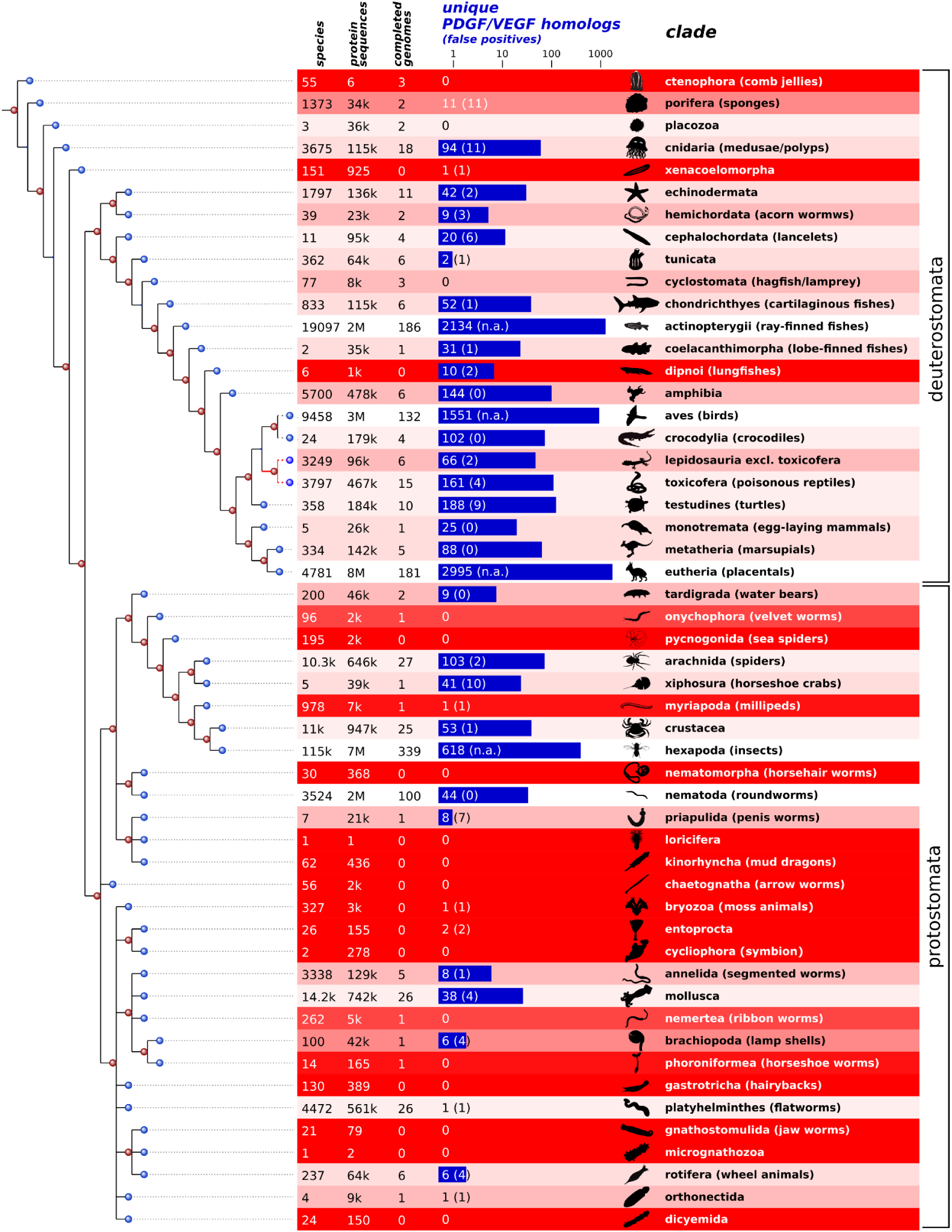
PDGFNEGF-like blast hits from 52 animal clades and their quantitative representation in the NCBI taxonomy and sequence databases. The number of blast hits from each clade is indicated in blue. The number of false positive hits is indicated in parentheses. When clades were represented by >500 species, false positives were not manually excluded (n.a.). The darker the red color, the less reliable the analysis results are due to the underrepresentation of the clade in the sequence databases. Most protostome phyla were underrepresented in the NCBI sequence databases. The current consensus tree of life [44] is shown on the left aligned with the clades.

Note that the relationship of Lepidosauria to Toxicofera, which might or might not be a clade, is shown with red lines. Not all 52 animal clades were equally well represented in the available sequence databases. In underrepresented clades, the absence of evidence for PDGF/VEGF-like genes was not taken as evidence of absence. To visualize this uncertainty, we used a heuristic formula to indicate the bias in the sampling, which takes into consideration the number of animal species of that clade in the NCBI taxonomy database, the number of sequenced genomes, and the total number of protein sequences for the clade available from the NCBI databases.

### PDGF/VEGF-like proteins of the least complex organisms resemble VEGF-C

The least complex animals where PDGF/VEGF-like proteins were identified are the Cnidaria (which include mostly medusae and corals). While 11 blast hits were from Porifera (sponges), which are less complex compared to Cnidaria, all of these were manually identified as false positives (five of these were genuine BR3P or BR3P-like proteins, see Supplementary Table 2). From the approximately 115,000 cnidarian protein sequences, 72 were identified as VEGF-like, including the previously described “VEGF” from the marine jellyfish *Podocoryne carnea* and fresh-water polyp *Hydra vulgaris* [45–47]. When we analyzed their amino acid sequences and compared them to modern-day VEGFs, they appeared more similar to the modern VEGF-C than to VEGF-A, -B, or PlGF. Similar to VEGF-C and VEGF-D, all but one of these contained the characteristic BR3P motif repeats C-terminally to the VEGF homology domain (VHD) (see Figure 2). Cnidarian VEGFs typically feature four BRP3 motif repeats after the VHD. Like *Drosophila melanogaster* PVF-2 and *C. elegans* PVF-1, they frequently lack one or both of the cysteine residues which form the intermolecular disulfide bonds in the mammalian PDGFs/VEGFs. When vertebrate PDGF/VEGF homologs are included in the generation of a phylogenetic tree, they cluster into one branch, indicating that PVFs likely originate from a single VEGF precursor gene in the genome of the most recent common ancestor of Vertebrata and Cnidaria (Figure 3). In seven out of the 18 gene-annotated Cnidaria genomes, VEGF-like sequences could be identified (*Acropora digitifera, Exaiptasia pallida, Hydra vulgaris, Nematostella vectensis, Orbicella faveolata, Pocillopora damicornis*, and *Stylophora pistillata*).

**Figure 2.**
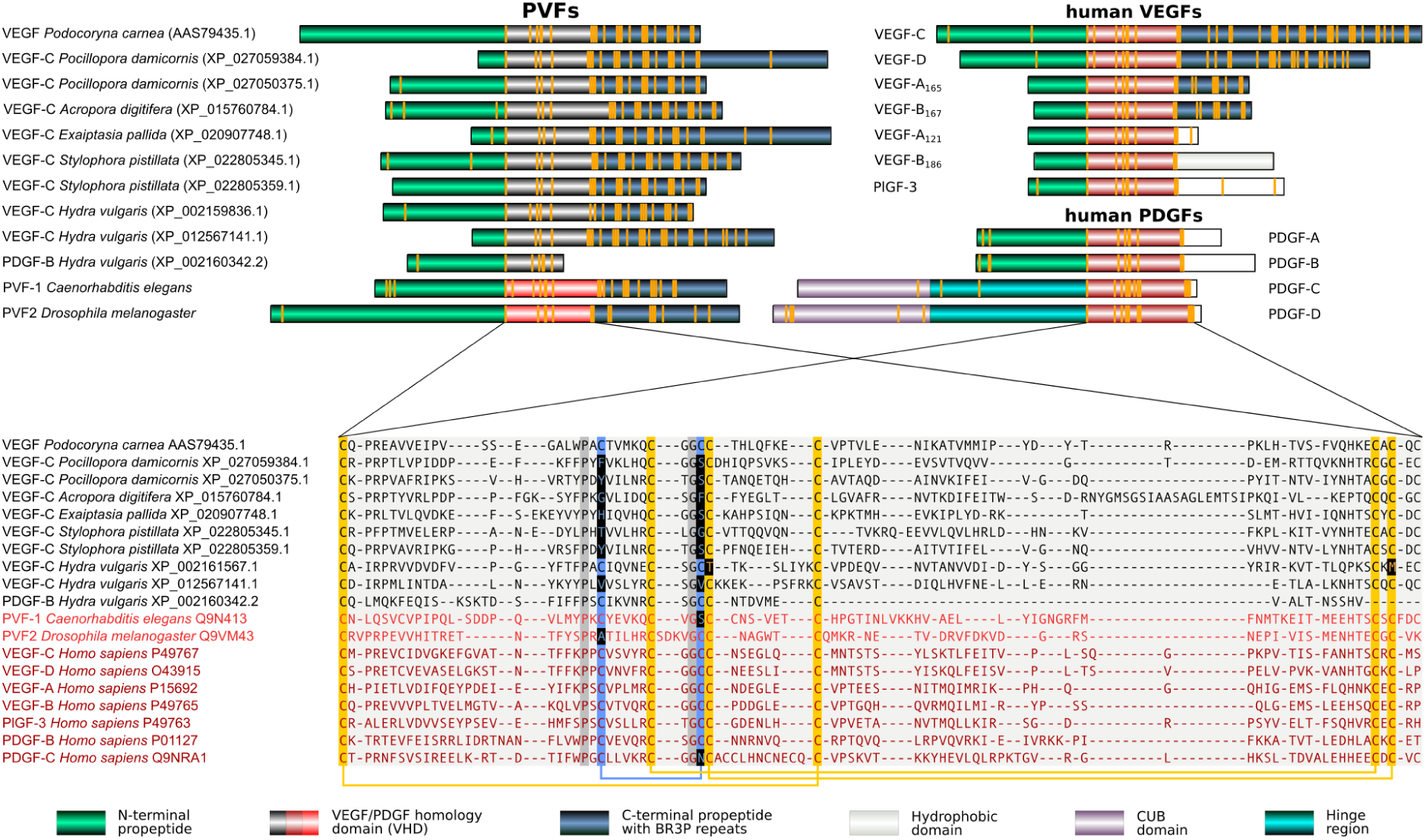
Comparison of the domain structure and alignment of the VEGF homology domain of Cnidaria and human PDGFs/VEGFs. 10 representative cnidarian VEGFs (sequences in black) were aligned with human VEGFs, selected PDGFs, *D. melanogaster* PVF2, and *C. elegans* PVF-1. Most cnidarian VEGFs show a typical cystine knot followed by three to five BR3P motifs similar to the human VEGF-C/D. BR3P motifs are completely absent from mammalian PDGFs and PlGFs, while the longer VEGF-A isoforms and the VEGF-B167 isoform contain one complete (CX10CXCXC) and one incomplete (CX_10_CXC) BR3P repeat C-terminally to the VEGF homology domain. In the alignment, the cysteines of the cystine knot are shown on an orange background, and the cysteines forming the intermolecular disulfide bridges are on blue background. The intermolecular disulfide bridges frequently appear absent in invertebrate VEGFs (marked by inverted coloring), but some of this might be an artifact of the alignment. 100% conserved non-cysteine residues are shown on dark grey background. The bridging pattern of the canonical PDGF/VEGF cysteines is indicated by connecting lines below the alignment.

**Figure 3.**
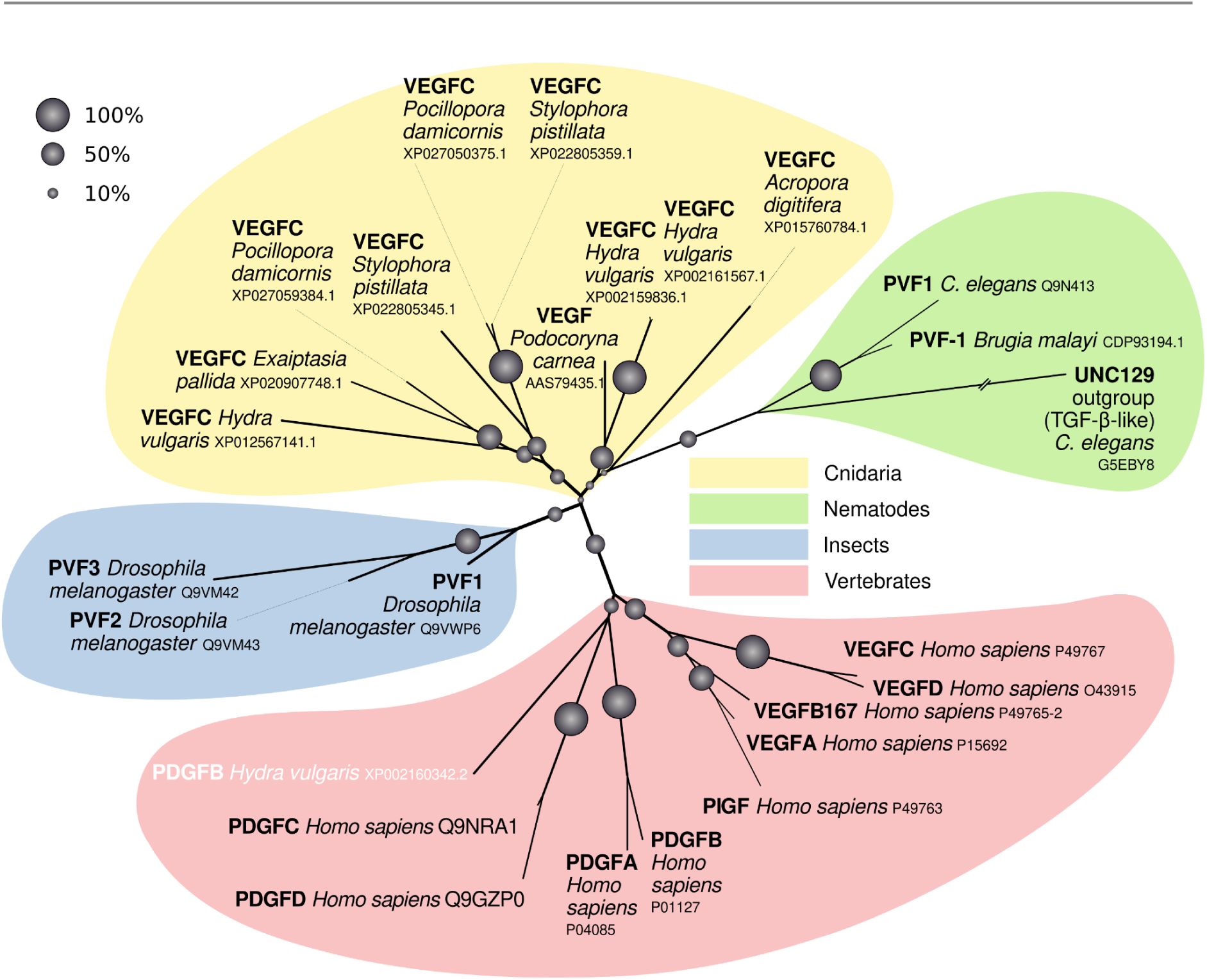
Vertebrate versus invertebrate VEGFs. A phylogenetic tree was calculated from an expanded set of sequences, aligning mostly the PDGF/VEGF homology domains. In this unrooted tree, *Podocoryna carnea* VEGF groups clearly together with all the other Cnidaria VEGF-C-like sequences, the exception being *Hydra vulgaris* PDGF-B (shown in white font color), which consistently groups with the mammalian PDGF-C/D group, with which it also shares other characteristics such as the near-complete lack of C-terminal sequences beyond the PDGF/VEGF homology domain. The confidence into branches is given as bootstrap values (% from 1000 repeats).

### Distribution of different VEGFs in the deuterostome branch

We did find VEGF-A-like proteins in Echinodermata, Cephalochordata, and Tunicata, all of which are clades in the deuterostome branch of the animal kingdom. However, all these animals - with the exception of the Tunicata - contained also VEGF-C-like proteins. Among all tunicate sequences, including the six completed tunicate genomes, only one PDGF/VEGF-like gene could be identified. The amino acid sequence of the predicted corresponding tunicate gene product showed a close homology to VEGF-A.

Not being a formal taxonomic group, there is considerable heterogeneity among fish. In bony fish (Osteichthyes), all five mammalian VEGFs (VEGF-A, PlGF, VEGF-B, VEGF-C, and VEGF-D) are ubiquitous. Previously, cartilaginous fish (Chondrichthyes: sharks, rays, skates, sawfish, and chimeras) were thought to lack PlGF and VEGF-B, but we could find in the majority of species sequences that showed the closest homology to PlGF and VEGF-B, respectively (Supplementary Figure S2). Only jawless fish (Cyclostomata: lamprey and hagfish) seem to be devoid of orthologs for PlGF, VEGF-B, and VEGF-D. When plotted along the branches of the phylogenetic tree of the animal kingdom, there is an overall expansion of VEGF diversity over time, while a few major branches undergo a collapse (Figure 4). The collapse coincides with a reduction in body plan complexity in the case of the Tunicata, which have either reduced the number of VEGF paralogs by eliminating VEGF-C-like sequences (*Ciona intestinalis*) or functional VEGF genes altogether (all other tunicates analyzed). However, a similar reduction of body plan complexity is not seen for the clade Archosauria with its extant members (birds and crocodiles), in which we did not find any signs of functional *VEGFB* genes. The same holds for Amphibia, in which we did not find functional genes coding for Placenta growth factor (*pgf*). While we also did not detect any *VEGFB* in Monotremata or any PDGF/VEGF-like proteins in Xenacoelomorpha, we did not consider these findings significant due to the incomplete sampling of these phyla in the sequence databases.

**Figure 4.**
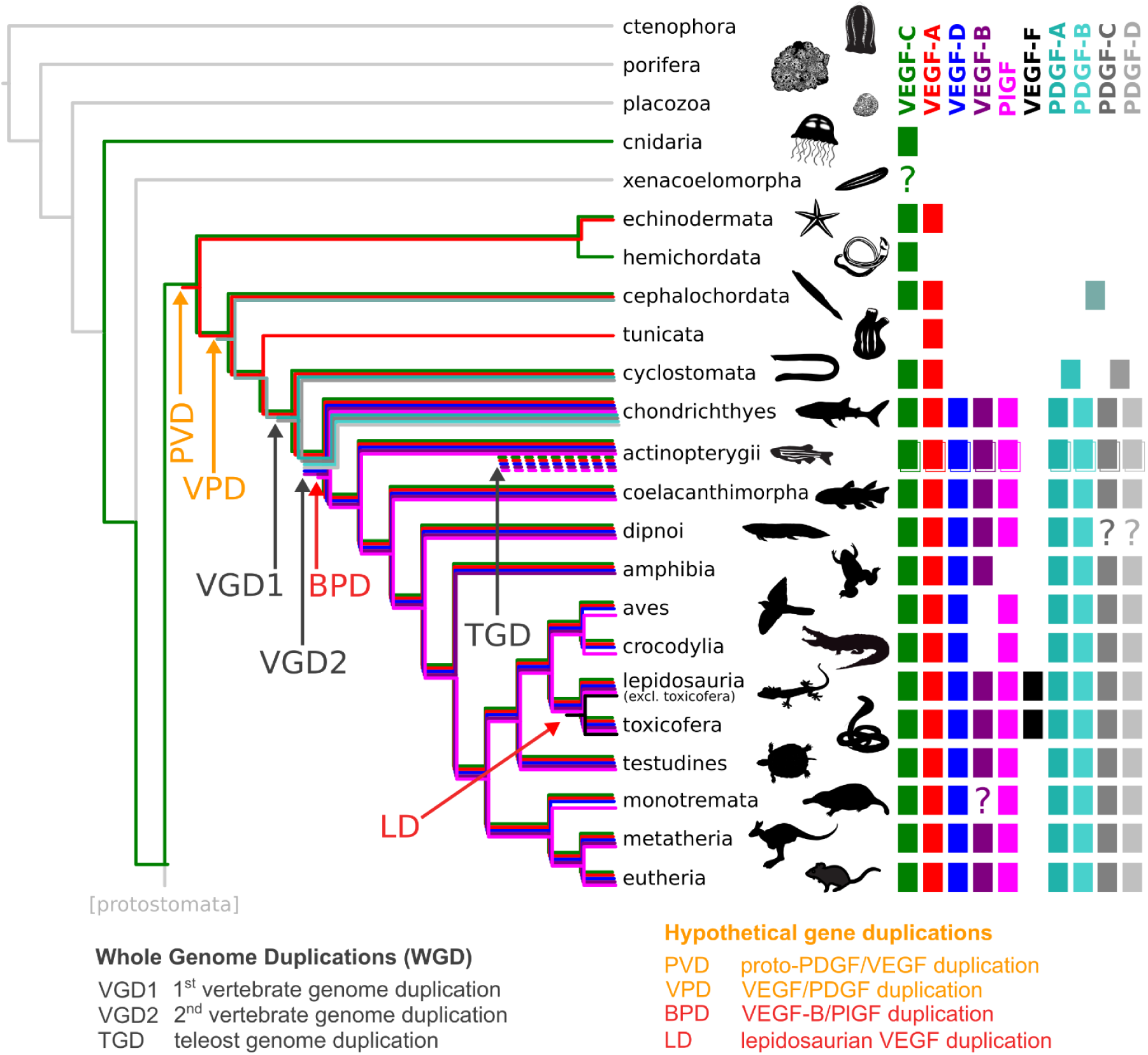
Occurrence of PDGF/VEGF genes in genomes of extant animal clades (excluding protostomes). Inferring from the occurrence of PDGFs/VEGFs in extant animal species, the first PDGF/VEGF-like protein appeared before the deuterostome/protostome split (DPS) and was likely most similar to the modern VEGF-C. The earliest expansion (proto-PDGF/VEGF duplication, PVD) appears after the DPS but prior to the known first vertebrate genome duplication (VGD1). One of the duplicated proto-PDGFs/VEGFs underwent partial removal of its C-terminal domain, resulting in a VEGF-A-like protein. The subsequent expansion of the family likely results from VGD1 and VGD2. Many mammalian genes have two orthologs in zebrafish that result from teleost genome duplication (TGD). Separate, limited gene duplication events (red arrows) explain the emergence of PlGF and VEGF-B soon after the VGD2-duplication (VEGF-B/PlGF duplication, BPD) and, more recently, of VEGF-F (Lepidosauria duplication, LD). Due to gene loss, some clades appear devoid of all or some VEGFs (VEGF-C: Tunicata, PlGF: Amphibia, VEGF-B: Aves, Crocodylia). In tunicates, the gene loss concurs with a massive reduction in morphological complexity, but not in amphibians, birds, and crocodiles. The separation of VEGFs and PDGFs (VEGF/PDGF duplication, VPD) likely happened soon after the first duplication of the proto-PDGF/VEGF as already Cephalochordata feature a single PDGF-like gene (see Supplementary Figure S2). Note that the absence of all VEGFs in Xenacoelomorpha, VEGF-B in Monotremata, and PDGF-C/D in Dipnoi could be due to the underrepresentation of these clades in the available sequence data. For reasons of clarity, the figure does not show a) the PDGFs lines on the branch leading to mammals starting from the VGD2 (since they are consistently present on that branch), b) the Salmonid Genome Duplication (SaGD), c) individual gene duplications not found in the majority of species of the tree branch (such as VEGF-C and PDGF-A duplications in some sharks and other fishes).

### Whole-genome duplications only partially explain the expansion of the PDGF/VEGF family

Whole-genome duplications (WGD) have contributed significantly to the increasing complexity of gene families and vertebrate evolution [48, 49]. When we overlayed the established WGDs and the emergence of novel PDGF/VEGF paralogs on the phylogenetic tree of the animal kingdom (Figure 4), the emergence of novel PDGF/VEGF family members coincides only partially with the proposed timing of the three WGDs. The first PDGF/VEGF (“Proto-PDGF/VEGF”) appears in the tree before the deuterostome/protostome split (DPS), but diversification is likely to have happened only after the DPS, since the protostome branch lacks the diversification pattern seen in the deuterostome branch. In the deuterostome branch, the first diversification happened likely at the protochordate stage, prior to the first vertebrate whole genome duplication (VGD1). PVFs from species in the protostome branch of the tree (insects, nematodes) do not show a clear diversification into distinct subgroups as seen in the vertebrate lineage, where PDGFs, angiogenic VEGFs (VEGF-A, -B, and PlGF), and lymphangiogenic VEGFs (VEGF-C and -D) have formed distinct subgroups. Already species that diverged soon after the DPS (Echinodermata) features two distinct VEGF homologs, which resemble the shorter, angiogenic VEGFs (VEGF-A-like) and longer, lymphangiogenic VEGFs (VEGF-C-like), followed soon by the separation of the PDGF branch, which becomes established before the divergence of the cephalochordate branch.

The VGD1 and VGD2 occurred within the direct line to mammals, while the third significant WGD event took place in the common ancestor of the teleost fish lineage [50], to which also zebrafish (*Danio rerio*) belongs. Although the VGD1 must have resulted in a duplication of the *proto-VEGFA* and *proto-VEGFC* genes, resulting in four VEGF homologs, we did not find any evidence that any of the duplicated *VEGF* genes permanently became established in the genome. Different from this, VGD1 is likely to have established the two PDGF subgroups by duplicating the *proto-PDGF* gene. However, a clear separation into a PDGF and a VEGF lineage had not occurred at this stage (see Supplementary Figure S2), and the proposed order of events is, therefore, the most parsimonious but not the only possible.

The VGD2 likely gave rise to the VEGF-C/VEGF-D subfamily by duplication of the *proto-VEGFC* gene, and to the VEGF-A/PlGF/VEGF-B subfamily by duplication of the *proto-VEGFA* gene. The emergence of the third member of the VEGF-A/PlGF/VEGF-B subfamily shortly after VGD2 and the emergence of VEGF-F are most parsimoniously explained by limited duplications in the common ancestor of all Actinopterygii and the common ancestor of all Lepidosauria, respectively.

### PDGFs and VEGFs in fishes

Because of recent discoveries in the developmental pathways of the fish vasculature [51], we wanted to know how successfully PDGF/VEGF ohnologs (WGD-generated homologs) withstood inactivation/pseudogenization, and whether PDGF/VEGF gene duplications can also be found in clades outside the teleost lineage. Thus, we analyzed RNAseq data from all fish species present in the FishPhylo database [52] (Figure 5). Despite significant heterogeneity, ohnologs of PDGFs/VEGFs could be identified in most other fishes, with the expected exception of the Holostei (bowfin and spotted gar). A notable exception in the teleost lineage is *vegfbb*, for which we did not find a single mRNA contig, and *pdgfba*, which seems to have been lost in five out of six salmonid species. Salmonids, on the other hand, show clear signs of having maintained some of their other PDGF/VEGF ohnologs originating from the Salmonid Genome Duplication (SaGD), most notably VEGF-A ohnologs.

**Figure 5.**
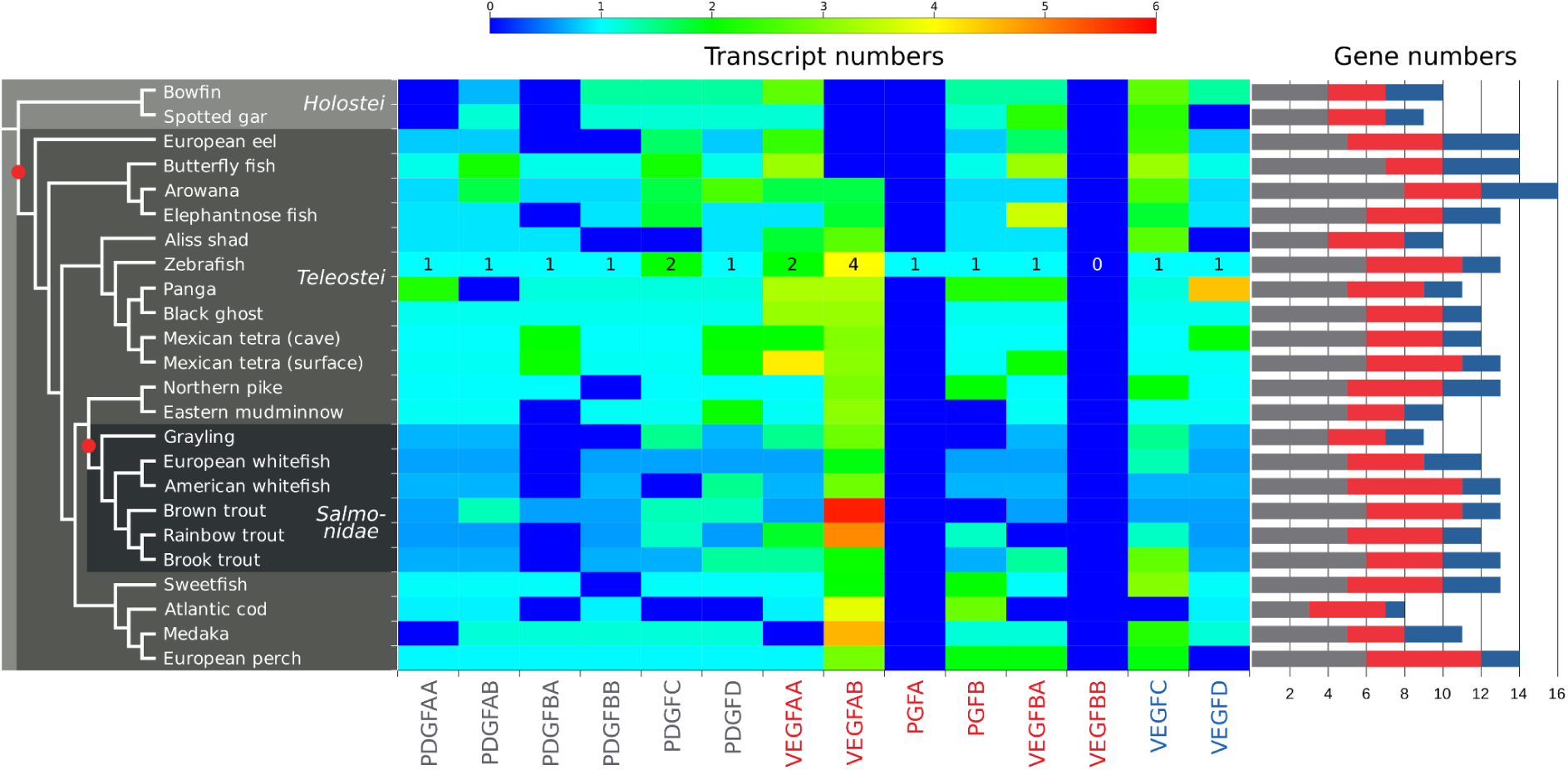
Numbers of different PDGF/VEGF mRNA transcript contigs and corresponding genes in 24 fish species as determined by RNA sequencing. While Holostei fishes did not undergo the Teleost Genome Duplication (TGD), they nevertheless feature individually duplicated genes such as *vegfc*. After the TGD, salmonids did undergo one additional round of genome duplication (Salmonid Genome Duplication, SaGD), and consequently, may feature - compared to humans - up to 4 times as many genes for each PDGF/VEGF, resulting theoretically in up to 36 different *pdgf/vegf* genes. About 20-50% of genes are functionally maintained after whole genome duplications due to neo- or subfunctionalization [53–56]; therefore, the expected number of functional *pdgf/vegf*-like genes in salmonids is between 12 and 20, which is in line with the number of genes deduced from the mRNA transcripts (Supplementary Tables 3 and 4). The heatmap has been normalized to zebrafish mRNA transcript numbers to compensate for the differences in the total number of mRNA transcript contigs obtained for each species. Whole genome duplications are shown as red dots on the cladogram on the left. Note that the transcript number is only weakly associated with the gene number.

We found VEGF-C duplicated in the fish clade Holostei, which diverged from teleosts before the teleost genome duplication. Only eight extant species comprise the extant Holostei lineage (the bowfin and seven gar species). All three fully sequenced genomes (*Amia calva*, *Atractosteus spatula*, and *Lepisosteus oculatus*) feature a duplicated *vegfc* gene indicating a single gene duplication early in the Holostei lineage (Figure 6A). When testing whether the *vegfc* genes in these fishes are still under purifying selection (i.e., whether gene inactivation or mutations are detrimental), we detected strong pervasive purifying selection throughout the coding region, with the strongest conservation in the receptor binding domain, followed by the silk homology domain (SHD) (Figure 6B). Among all fish species, individual gene duplications were found frequently for VEGF-C but occasionally also observed for other genes such as VEGF-A (Figure 6C).

**Figure 6.**
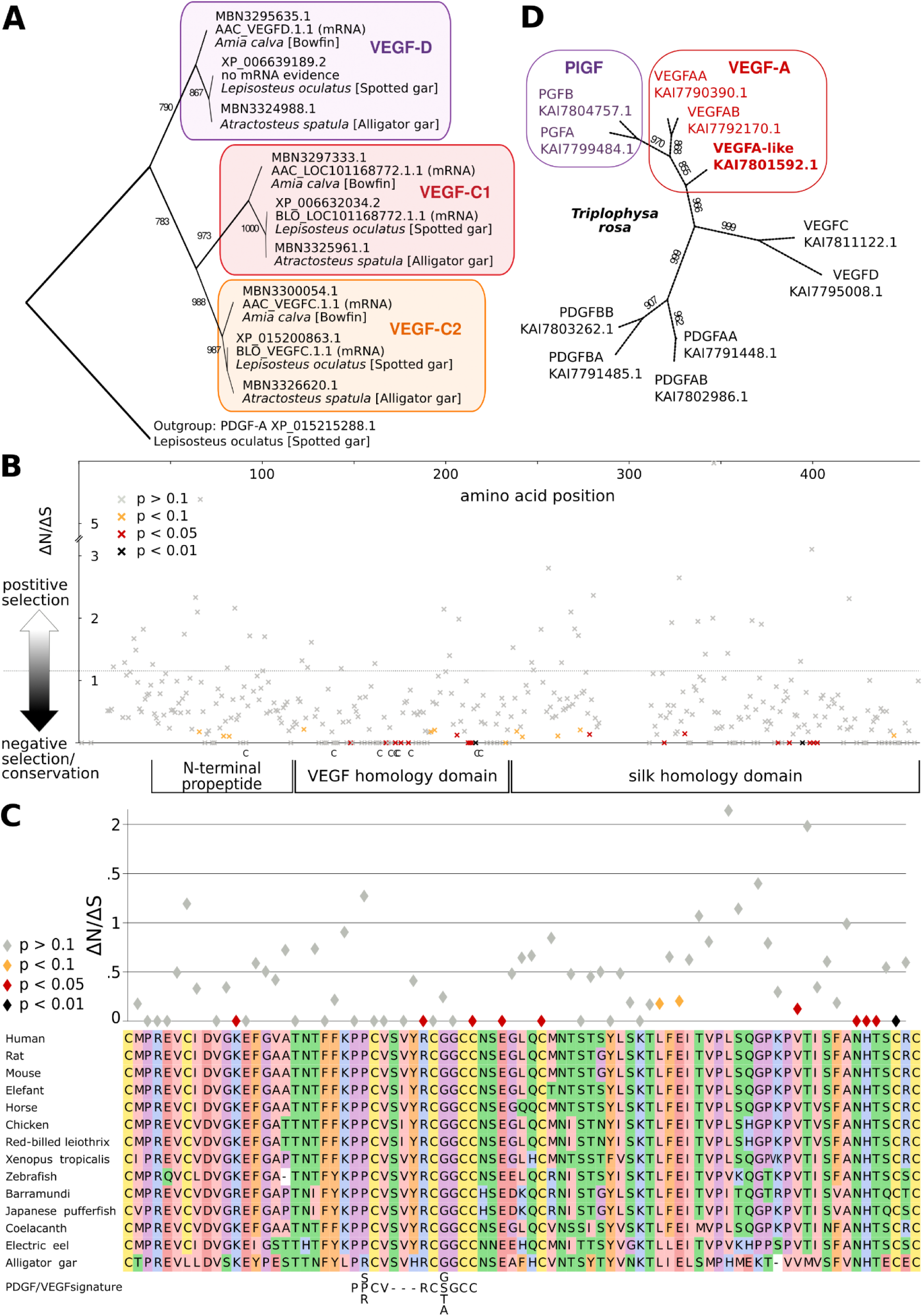
Individual gene duplications contribute to VEGF diversity in fishes. (A) The *vegfc* gene is duplicated in all Holostei species, and for the Bowfin and the Spotted gar, there is mRNA evidence for gene expression of both paralogs, while there was no evidence for the expression of *vegfd* in the Spotted gar (see also Supplementary Table 4). (B) Holostei *vegfd* and *vegfc* genes are under purifying selection, most notably in the VEGF homology (VHD) and silk homology domain, where there appears to be strong pressure to maintain the cystine-knot structure. Cysteine residues of the VHD are indicated below the x-axis. (C) Detailed view on the conservation of the VHD, aligned with representative VEGF-C amino acid sequences and the PDGF/VEGF signature. Please note that sites without synonymous codon changes (ΔS = 0) are not displayed. (D) *vegfa* is also sometimes individually duplicated in some fish species, as shown for *Triplophysa rosa*.

### PDGFs/VEGFs are not absent in jawless fishes but might be absent in parasitic Cnidaria

As shown in Figures 5 and 6, individual gene duplications or gene losses were not uncommon in fishes, but a complete apparent absence of PDGFs/VEGFs was initially seen in the extant jawless fishes (Cyclostomata), sea lamprey (*Patromyzon marinus*) and inshore hagfish (*Eptatretus burgeri*). Based on our previous results, we reasoned that the lack of cyclostomate PDGF/VEGFs in protein databases must be an artifact, perhaps due to a failure of gene prediction or annotation. In fact, a manual inspection of the Ensemble [57] lamprey and hagfish genomes showed - as predicted by our phylogenetic tree - four PDGF/VEGF-like sequences, and an improved version of the Ensemble genebuild pipeline added the above proteins to the Uniprot database in 2022 (see Supplementary Figure S2). However, the inshore hagfish sequences have not yet been migrated to the (blastable) NCBI protein database. The occurrence of four PDGF/VEGF-like genes in Cyclostomata supports the Early-1R hypothesis. The occurrence of four PDGF/VEGF-like genes in Cyclostomata supports the Early-1R hypothesis.

Since we had identified multiple PDGF/VEGF-like proteins in seven Cnidaria species, we reasoned that the lack of these in other Cnidarians could also be an artifact. For in-depth inspection, we chose *Thelohanellus kitauei*, which is a genome-sequenced cnidarian represented with 14792 protein sequences in the Uniprot database. We could not identify genes coding for PDGF/VEGF-like proteins in its genome using very relaxed degenerate search criteria. *T. kitauei* belongs to the endoparasitic myxozoa branch of Cnidaria, which is characterized by a reduction in genome size and gene depletion [58], likely explaining the lack of PDGF/VEGF-like sequences in this and other parasitic Cnidaria.

### Both VGD1 and VGD2 contribute to PDGF expansion

Kipryushina et al. [59] place the emergence of PDGFs after the divergence of Echinoderms and Chordates, notwithstanding early reports of PDGF/PDGFR signaling in sea urchins [60]. Concurring with Kipryushina, the PDGF/VEGF-like proteins identified from the known 11 sea urchin genomes are highly homologous to the proto-VEGF-C that we found in Cnidaria, and we found the first PDGF-like growth factors in Cephalochordata (lancelets). Lancelet genomes feature one or two PDGF-like growth factor genes and one VEGF-C-like gene (see Supplementary Figure S2). However, only VEGF-C-like growth factors can confidently be placed into the corresponding ortholog group due to their BR3P signature. Most tree topologies placed the other cephalochordate PDGF/VEGF family members between VEGF-Cs and PDGFs, albeit with a rather low bootstrap support values. The cephalochordate PDGFs/VEGFs were likely duplicated by VGD1 since Cyclostomata (hagfish, lamprey) already feature four PDGF/VEGF genes. While the VGD2 explains the emergence of PDGF-D by duplicating the proto-PDGF-C/D, the exact order of the preceding events in the early VEGF/PDGF evolution cannot be reconstructed with confidence as the support values of the phylogenetic tree reconstructions remained low, independently of the methods used (see Supplemental Figure S2). This is not surprising if assuming a monophyletic origin of Cyclostomata [61, 62] but results in the fact that the sequences from Cyclostomata are not very informative about the early PDGF/VEGF evolution.

### VEGF-F can be found in several Lepidosauria, not only in venomous snakes

In 1999, it was recognized that the hypotensive factor from the venom of *Vipera aspis* [21] is a VEGF-like molecule [22]. Our analysis shows that VEGF-F is not limited to venomous snakes but is also found in non-venomous snakes (e.g., *Python bivittatus*, XP_025024072.1) and lizards, independently of whether they are venomous or not (see Supplemental Figure S3). Amino acid sequence alignments of VEGF-Fs show a similar high homology to both VEGF-A and PlGF, and based on phylogenetic trees, it appears likely that either VEGF-A or PlGF served as an evolutionary template for VEGF-F. Because we identified VEGF-F orthologs also in lizards, e.g., in the common wall lizard (*Podarcis muralis*, XP_028597744.1) or the gekko (*Gekko japonicus*, XP_015284783.1), the gene duplication likely happened early in the Lepidosauria lineage, before the invention of venom. However, we found a clear subdivision in the VEGF-F branch between “viper” VEGF-Fs and “non-viper” VEGF-Fs. Only for the VEGF-Fs from the viper branch, the venom character of the VEGF-Fs has been experimentally verified.

### Viral VEGFs

Another biological entity that has co-opted PDGFs and VEGFs for its own purposes are viruses. To analyze the relationship between all available viral VEGF sequences (VEGF-Es), we constructed a phylogenetic tree of all VEGF-E sequences that were identified in the main analysis (see Figure 7 for the simplified tree and Supplementary Figure S4 for the complete tree). Sequences coding for VEGF-like genes were found in the genomes of at least four different virus clades: orf virus (ORFV), pseudocowpoxvirus (PCPV), bovine pustular stomatitis virus (BPSV), and megalocytivirus (MCV). ORFV, PCPV, and BPSV are collectively known to infect at least ten mammalian species, while MCVs have been detected in at least eight fish species. The most parsimonious phylogenetic trees suggest that VEGF-Es originate from VEGF-A. We did not find any evidence of recent multiple host-to-virus gene transfers as all vertebrate VEGFs formed tight clusters, which were well separated from the VEGF-E clusters. We did not observe a separation of the host species with the phylogeny. However, only the ORFV cluster probably contains enough sequences to allow for any such separation to become apparent.

**Figure 7.**
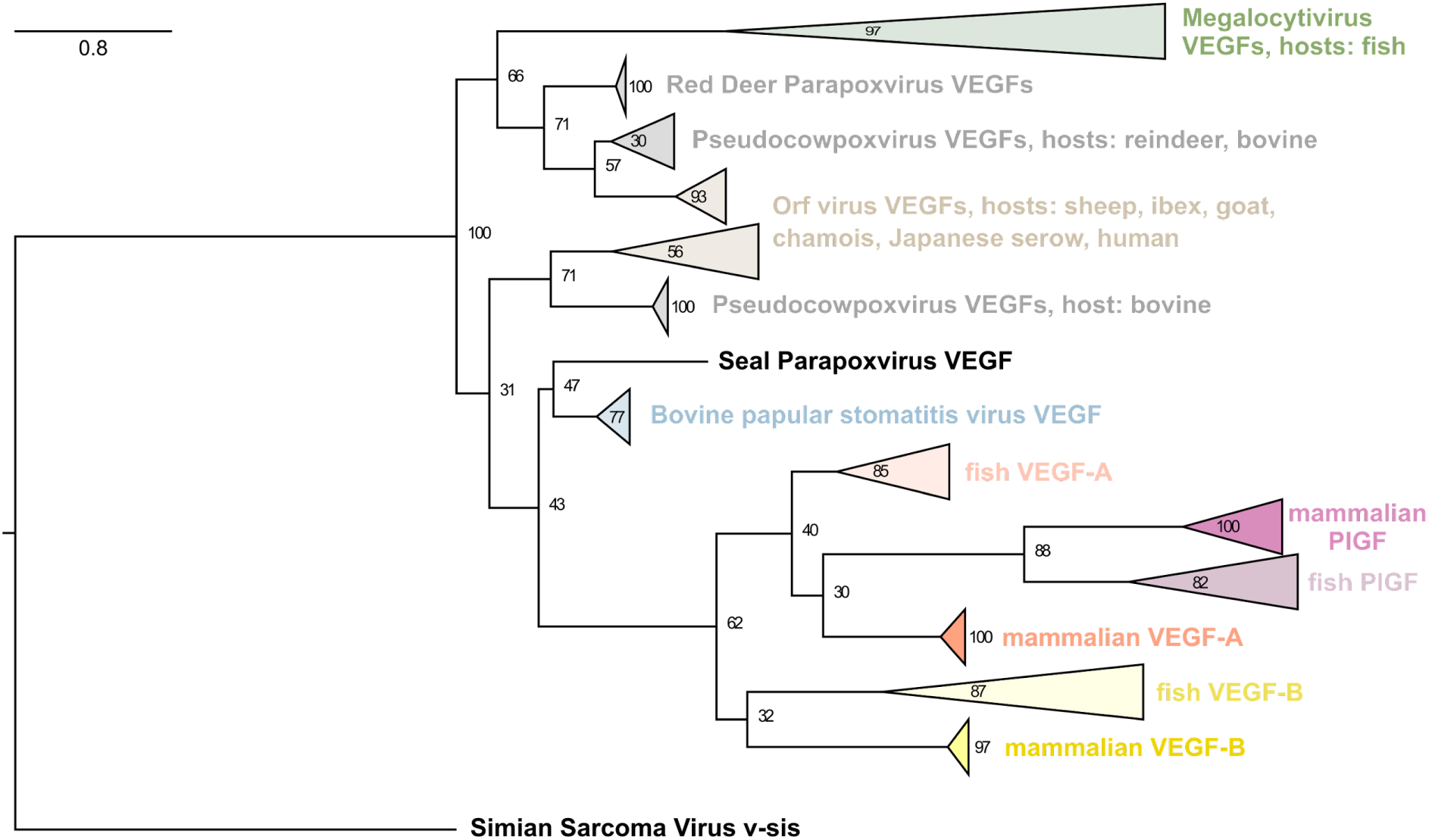
Phylogenetic protein tree of VEGF-E. Without exception, all VEGF-E and all non-viral VEGF sequences cluster together, arguing that none of the known VEGF-Es originates from recent host-to-virus gene transfer events. The protein tree is compatible with a single origin of all VEGF-E, but due to the significant distance, convergent evolution cannot be excluded. Based on the VEGF sequence alone, assignment to the Pseudocowpoxvirus or Orf virus group is not possible, since several VEGF sequences derived from different viruses are identical (e.g., reindeer PCPV VEGF is identical to the VEGF sequence from the PCPV reference genome VR634). In this branch of the tree, cross-species transmissions have been reported, including to humans [63]. The expanded tree with all leaves is shown in Supplementary Figure S4.

**Figure 8.**
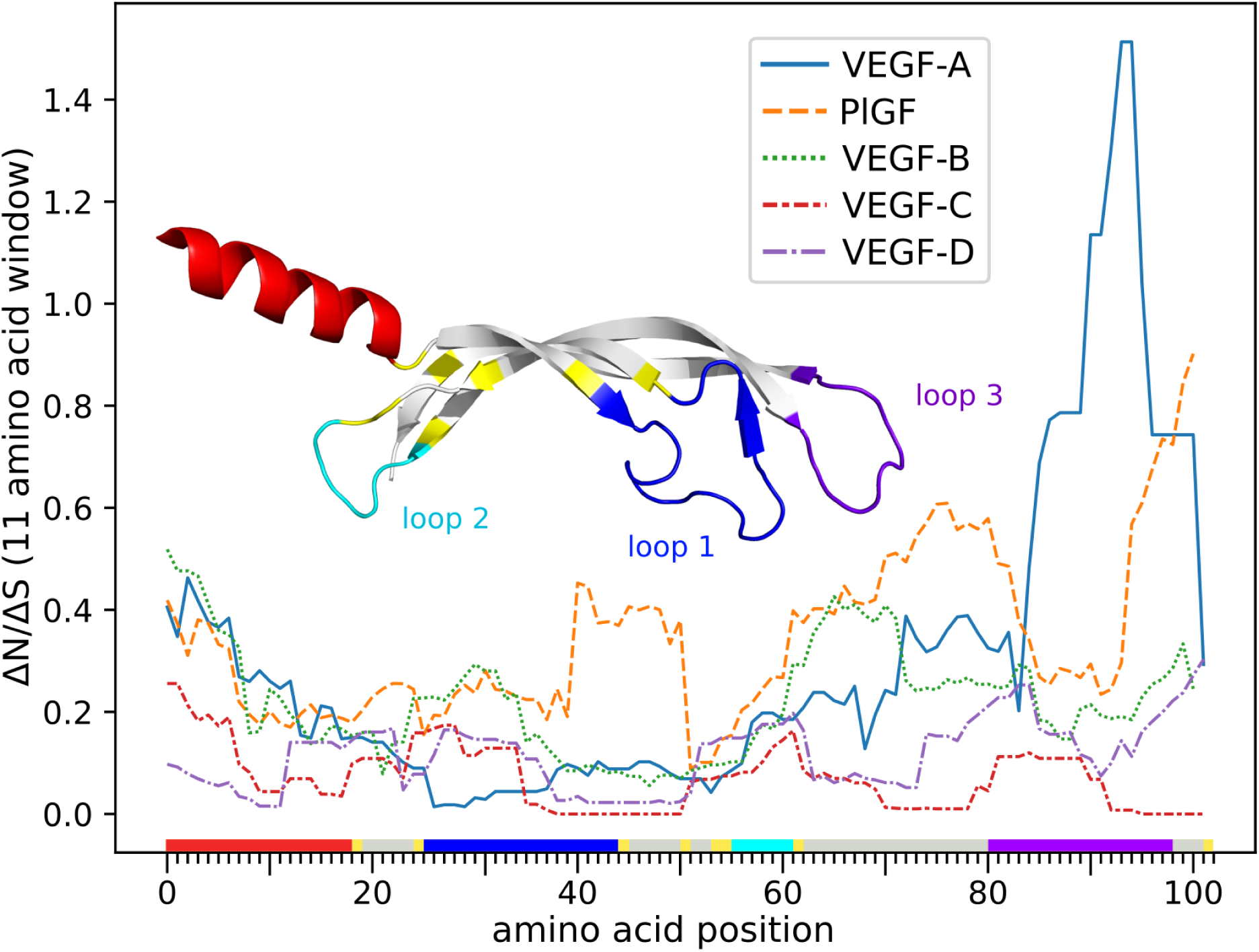
Conservation of the VEGF homology domain. The ratio of non-synonymous to synonymous mutations was inferred for all sites over the whole tree and averaged over an 11-amino acid window. VEGF-C shows the highest conservation, while the evolutionary younger family members appear to be more volatile. The amino acid position on the x-axis is color-coded to correspond to the location of the residue in the 3D structure of the monomeric VEGF-C protein [65]. Note that the peak at the end of the VEGF-A sequence might or might not be an artifact resulting from the difficulty of generating a good alignment for loop 3 (shown in purple). Alternatively, the peak might correspond to adaptations in the receptor interaction, as loop3 carries major determinants for receptor binding [66, 67].

### PDGF/VEGF-like molecules of the protostome branch

The sequence coverage of the protostome branch of the animal kingdom was much weaker than that of the deuterostome branch. For 16 out of 29 invertebrate phyla, we could not find a single genomic draft sequence, and almost all available genomic data covered only six phyla: flatworms, roundworms (nematodes), molluscs, crustaceans, insects, and spiders. While nematodes and flatworms do not possess blood circulation, insects, spiders, and crustaceans feature a so-called “open circulation”, where the hemolymph is circulated inside a body cavity (hemocoel). PDGF/VEGF-like molecules were identified in all these six phyla except for flatworms. Less than half of the 100 completed nematode genomes contained PDGF/VEGF-like genes. Among these are notably many *Caenorhabditis* PVFs, like *C. elegans* PVF-1 (NP_497461.1), but the majority were found from parasitic nematodes, including intestinal parasites like *Ancylostoma duodenale* (KIH56282.1), lymphatic parasites like *Brugia malayi* (CDP93194.1) and *Wuchereria bancrofti* (VDM19972), and conjunctival parasites like *Loa loa* (XP_003142823.1). PDGF/VEGF-like proteins appear common if not pervasive among molluscs, crustaceans, insects, and spiders, and can be identified in most genome-sequenced species.

### Conservation of the VEGF sequences

Except for VEGF-A and VEGF-C, the physiological and pathophysiological role of VEGF family members is still under debate. Some of them (*Vegfb* and *Vegfd*) can be deleted in mice without any major phenotype [12, 13, 64]. To compare the evolutionary pressures acting on different VEGFs, we analyzed the conservation of their coding sequences. We compared nonsynonymous and synonymous substitution rates inferred by a maximum-likelihood approach. The VEGF-C sequence was most strongly conserved, followed by VEGF-D, while PlGF showed the highest variability while still being relatively conserved. VEGF-A was overall also very conserved but showed a peak of variability at the very end of the VHD corresponding to loop 3, which is a major carrier of the receptor binding epitopes for VEGF.

### VEGF-A splice isoforms and diversity quantification

Splice isoforms are a means of generating diversity at the protein level from a single gene. Within the PDGF/VEGF family, alternative mRNA splicing is very unequally used to generate diversity: the hemangiogenic VEGFs (VEGF-A, PlGF, and VEGF-B) all feature at least two major splice isoforms, while the lymphangiogenic VEGFs (VEGF-C and VEGF-D) seem to generate all of their protein diversity post-translationally by alternative proteolytic processing [68]. The <u>Uniprot protein entry for human VEGF-A</u> describes no less than 17 splice isoforms, eight of them with evidence at the protein level. When reimplementing the DIVAA software [69] in Biopython to quantify the diversity of VEGF sequences, we realized that for genes rich in splice isoforms such as VEGF-A, an accurate assignment of isoforms is paramount since indels are not handled well by alignment and tree-building algorithms. Using the length of the coding region and comparisons with a reference list of known VEGF-A mRNA isoforms, we programmatically sorted VEGF-A protein sequences into four major buckets, corresponding to the human 121-, 165-, 189-, and 206-isoforms. When we counted the number of sequences for these four VEGF-A isoforms, we found a ratio of roughly 9:4:3:1 for the 189, 165, 121, and 206 isoforms, indicating that VEGF-A_189_ might be the predominant VEGF-A isoform in many species.

## Discussion

In this study, we have identified homologs of PDGFs/VEGFs in most animal phyla that show tissue organization (i.e., excluding sponges) and for which more than a few genome assemblies and gene predictions exist. Despite their pervasive occurrence in many branches of deuterostomes and protostomes, the data clearly support the notion that some animal phyla are completely or partially devoid of PDGF/VEGF-like molecules, and this might, above all, apply to clades with secondarily reduced body plans like Tunicata, or for the phyla Xenacoelomorpha or Dicyemida. Especially for many of the protostome phyla, not much genomic or mRNA data is available. Genome assemblies are often lacking, and the available prediction algorithms might not be very reliable as these animals are rarely the subject of genomic research. For these phyla, the lack of PDGF/VEGF-like proteins is a provisional hypothesis.

After hypothetical proteins are predicted from genomic sequences, programmatic bioinformatics workflows typically assign them to homology groups (e.g., using PANTHER [70]), resulting in automatic annotation like <u>PREDICTED, VEGF-C</u>. Despite this approach, many PDGF/VEGF homologs fail to be programmatically categorized into one of the 10 ortholog groups (<u>VEGF-A</u>, <u>PlGF</u>, VEGF-B, <u>VEGF-C</u>, VEGF-D, VEGF-F, PDGF-A, PDGF-B, PDGF-C, PDGF-D). Our algorithm emulates crowd-sourcing by comparing uncategorized homologs to the most closely related manually and programmatically annotated PDGFs/VEGFs and establishes a majority opinion, allowing for the categorization of the majority of uncategorized vertebrate proteins into one of the ortholog groups. This crowd-sourcing combines human annotation of gene and protein records with the tree-building and clustering methodology used for the protein trees available at Ensemble (https://m.ensembl.org/info/genome/compara/homology_method.html), which are used for the automatic annotation.

Based on the phylogenetic tree of the animal kingdom and our analysis of PDGF/VEGF homologs in different animal clades, the emergence of the earliest PDGF/VEGF-like molecule (“proto-PDGF/VEGF”) predates the establishment of the bilaterian body plan [71] and the split of the animal kingdom into deuterostome and protostome organisms before the start of the Cambrian about 540 MYA [72]. Intriguingly, this “proto-PDGF/VEGF” most likely featured a domain structure characteristic for the modern lymphangiogenic VEGF-C/VEGF-D subclass having long N- and C-terminal extensions flanking the VHD and a characteristic repetitive cysteine residue pattern (the BR3P repeat) at its C-terminus. Concurrent with the relatively rapid evolution of different body plans during what has been termed the “Cambrian Explosion”, the proto-PDGF/VEGF undergoes diversification establishing a VEGF-A-like and a VEGF-C-like branch and spinning off the PDGF branch. The deuterostome/protostome split predates both the VEGF diversification and the PDGF spinoff, which explains the difficulty of classifying PDGF/VEGF-like molecules on the protostome branch (“*Drosophila* VEGFs”, “*C. elegans* VEGF-C”) as either PDGFs or VEGFs.

In the protostome branch, we can detect PDGF/VEGF-like factors (PVFs) in all but one clade for which substantial sequencing data is available. All 26 genome-sequenced flatworms seem to get along without any PVFs. Contrary to this, insects, mollusks, and segmented worms (Annelida) often feature more than one *Pvf* gene, whereas most nematodes feature only one. *C. elegans* PVF-1 is remarkable in that it is still - after more than 500 MYA of evolutionary separation - able to activate the human VEGF receptors-1 and -2 [38]. Protostome animals do not feature a cardiovascular system, with exceptions among mollusks and segmented worms (Annelida). The phylum Annelida contains species such as the earthworm *Lumbricus terrestris*, which features a closed circulatory system displaying vascular specialization and hierarchy, including a vessel-like heart with valves, larger blood vessels, capillaries, blood containing free hemoglobin, and a renal filtration system [73]. Until now, no protostome PVF has been shown to play any role in vascular development, and a unifying picture of PVF function in invertebrates has yet to emerge.

With PlGF, VEGF-B, VEGF-D, and VEGF-F, further specializations happen in both the VEGF-A and the VEGF-C lineage after the Cambrian period in the deuterostome branch. Instrumental for this specialization is most likely the VGD2, which doubled the number of PDGF/VEGF-like genes. When ignoring teleost fishes, whole genome duplications are responsible for about half of the newly emerging PDGFs/VEGFs, the other half requiring duplication events at the gene or chromosome level (Figure 4).

Echinodermata are the most simple animals that display PDGF/VEGF specialization at the gene level. The least complicated explanation for the existence of both VEGF-A-like and VEGF-C-like proteins in Echinodermata is that the first gene duplication of the proto-PDGF/VEGF happened prior to the VGD1. For the same reason, the separation of the PDGF lineage also likely predates the VGD1, resulting in PDGFs being present in cephalochordates (lancelets), which are the most simple organisms to have a pressurized vascular system in which the blood is moved around by peristaltic pressure waves created by contractile vessels [74]. In line with this notion is the important role of PDGFs in the supportive layers that stabilize blood vessels (pericytes, smooth muscle cells) [75].

Despite the availability of genome assemblies from six different tunicate species, the programmatic approach identified only one VEGF-like molecule in tunicates, *Ciona intestinalis*. This was surprising since there is prior data indicating the role of VEGF/VEGFR signaling in the circulatory system of tunicates [76]. However, the relationship between the receptor tyrosine kinase that was cloned from the tunicate *Botryllus schlosseri* and PDGF receptors, FGF receptors, and c-kit is not clear. The immunological detection used antibodies directed against human VEGFRs or VEGF-A, and the TK inhibitor PTK787 might have as well inhibited PDGF receptors and c-kit. Although the *C. intestinalis* VEGF-like growth factor appears most similar to VEGF-A, its phylogeny also allows descendence from the PDGF or VEGF-C branch. In that case, the similarity to VEGF-A might have originated from convergent evolution (“long branch attraction”). Since none of the other tunicate species seems to feature PDGF/VEGF homologs, horizontal gene transfer could be an alternative explanation.

### Absence of PlGF and VEGF-B from entire animal vertebrate classes, VEGF-F more common than expected

Most striking was the absence of some VEGF family members from entire animal classes. We did not find any PlGF ortholog in Amphibia and also no VEGF-B ortholog in the clade Archosauria, which includes extant birds and crocodiles as well as extinct dinosaurs. Since bony fish feature both PlGF and VEGF-B, the absence of VEGF-B in extant Archosauria and PlGF in extant Amphibia likely represents an example of lineage-specific gene loss. While five avian protein sequences are annotated as “VEGF-B” or “VEGF-B-like” in the searched database, they did separate on a phylogenetic tree to the same branch as the VEGF-A sequences (data not shown). In addition, their genes did not show the typical exon-intron structure, which is characteristic of VEGF-B with overlapping open reading frames leading to two different protein sequences due to a frameshift [77].

As a counterpoint to the missing PlGF and VEGF-B, we observed, to our surprise, that VEGF-F is more common than generally thought. Discovered as a venom compound of vipers [21, 22], it was initially thought to be limited to venomous reptiles. However, we did detect VEGF-F-like sequences in non-venomous lizards and gekkos. VEGF-F is more or less pervasive throughout large parts of the lepidosaurian lineage, with occurrences in species so diverse as the Green anole and the Japanese gekko (which are located distantly from each other on the lepidosauran tree). For this reason, VEGF-F likely evolved early on in the evolution of Lepidosauria prior to the invention of venom (Supplementary Figure S3). At this moment, it is unclear which functions VEGF-F might have originally fulfilled before it became co-opted as an integral viper venom component. In vipers, VEGF-F expression is highly restricted to the venom glands [78], where it acts by accelerating venom spread by inducing vascular permeability and by incapacitating the prey by lowering blood pressure. However, it is conceivable that VEGF-F still fulfills its original, non-venom function in the non-viper branch of the VEGF-F tree.

### Complete absence of PDGFs/VEGFs

The absence of individual PDGFs/VEGFs from a species’ proteome can be either real or only apparent due to incomplete sampling or an artifact of the bioinformatics analysis pipeline. We generally found very few exceptions to the clade-specific pattern of PDGF/VEGF occurrence in terrestrial vertebrates, which all featured the same set of PDGF/VEGF paralogs, confirming the reliability of the respective genome sequencing and gene prediction pipelines. In our programmatic screen, PDGF/VEGF-like sequences were apparently completely absent in some clades for two different reasons:

1. A lack of data (false negatives): PDGFs/VEGFs were apparently absent from clades where there was no comprehensive genomic data or the genomic data had not been analyzed (e.g., sea spiders or velvet worms).
2. A true absence: PDGFs/VEGFs were absent from clades that are likely truly devoid of VEGF-like molecules (e.g., flatworms, where a substantial number of genomes have been sequenced and analyzed).

With increasing sequencing coverage, false negatives will disappear as has happened for Cyclostomata during the writing of this manuscript. The occurrence of four PDGF/VEGF-like genes in Cyclostomata supports the currently largely accepted Early-1R hypothesis (i.e., that the VGD1 happened before the divergence of Cyclostomata). While recent data suggest an early hexaploidization event for the Cyclostomata branch [79], we did not find evidence for more than four PDGF/VEGF genes in any cyclostomate genome.

### Viral VEGFs

While many viruses indirectly induce angiogenesis [80], some viruses encode their own VEGF homologs. These proteins have been collectively termed “VEGF-E”. Viral VEGFs have been reported from parapoxviruses, which cause skin lesions in their respective mammalian hosts [81–83]. In these viruses, the VEGF-E gene is specifically responsible for swelling and vascular proliferation [84]. Based on the sequence homology to VEGF-A, VEGF-E is believed to have been captured from a host during viral evolution [82], similar to the v-sis oncogene, which is believed to be derived from captured host PDGF-B sequences [25]. Our database search confirms that viral VEGF homologs exist not only in four species of the parapoxvirus genus but also in the very distantly related megalocytiviruses, which infect fish [85]. Surprisingly, despite their non-overlapping host range, both the fish and mammalian viral VEGF-Es might originate from one single acquisition from a mammalian host. Unlike megalocytiviruses, which infect fish (and occasionally amphibians), parapoxviruses have a very broad mammalian host range, which occasionally includes humans but is mostly covering domesticated and wild ungulates [86, 87]. While parapoxvirus infections are typically self-limiting, megalocytiviruses cause considerable economic damage to aquaculture. The pathophysiology of megalocytiviral diseases is not well understood. The infection leads to perivascular cell hypertrophy [88], and VEGF-E might facilitate virus dissemination via increasing vascular permeability.

### The “silk homology” domain

The alignment of the accessory domains of VEGFs is non-trivial since these domains contain a variable number of repetitive motifs and thus, the evolutionary history (i.e., which repeats got lost or added in which order) is perhaps impossible to deduce with reasonable accuracy. This holds true most notably for the SHD of VEGF-C and VEGF-D, which consists of several complete and incomplete Balbiani ring-3 protein (BR3P) repeats. The C-terminal tails of the VEGF-A_165_ and VEGF-B_167_ isoforms show a reduced set of these BR3P repeats, and in the case of VEGF-A, this domain has been named “heparin binding domain” (HBD). “Heparin binding” (i.e., binding to the extracellular matrix and cell surfaces) is one function of the SHD [89], and the HBDs of VEGF-A_165_ and VEGF-B_167_ have developed a stronger heparin affinity compared to VEGF-C or VEGF-D, which perhaps allowed for the reduced size of the HBD compared to the SHD of VEGF-C. In addition, the SHD of VEGF-C is required to keep VEGF-C inactive after secretion, likely by sterical hindrance [90], which is not a requirement for the longer VEGF-A isoforms, where inactivity might be mediated by sequestration mediated by the HBD [91]. However, when ECM-bound VEGF-A_189_ or VEGF-A_206_ are in direct contact with endothelial cells, some signaling seems to be possible [92]. Also, the VEGF-A_165_ isoform, which ranges in its ECM association strength between the soluble VEGF-A_121_ and the strongly ECM-bound VEGF-A_189_ and VEGF-A_206_ isoforms, can signal while being ECM-associated. However, the signals appear distinct from free VEGF-A_165_ [93, 94]. Our analysis shows that the SHD was likely an essential part of the proto-PDGF/VEGF. Given that the ECM-binding SHD is larger than the receptor-activating VHD, it is surprising that it has been maintained over several hundreds of million years. Since proteins are typically kept inactive by much shorter propeptides, it is tempting to speculate that the SHD must have some additional function beyond keeping VEGF-C inactive. One potentially important function could be the establishment of a VEGF-C gradient.

VEGF-A gradients are instrumental in the patterning of vascular networks [95–97]. They are believed to result from the interaction of the heparin-binding, longer VEGF-A isoforms with extracellular matrix and cell surface heparan sulfate proteoglycans and to be essential for directed vascularization during embryogenesis [98, 99]. Although the VEGF-A_165_ isoform is considered the major isoform in humans [100], the stronger ECM-binding VEGF-A_189_ (and other long isoforms characterized by recruiting an additional exon from the intron 5-6 compared to VEGF-A_165_) might be the predominant isoforms in many other animals, as they dominate in sequence databases. However, an increasing number of protein database entries are predictions from genome sequencing projects. Hence it is difficult to decide whether this abundance of the 189 isoforms reflects true mRNA dominance in most organisms or whether it results from a greedy bias in the splice prediction algorithm favoring longer isoforms. Supporting the hypothesis that in many animals, the longer VEGF-A isoforms are dominant is the inability to computationally map isoforms corresponding to the human VEGF-A_165_ (and sometimes also VEGF-A_121_) from genomic data in many animals, including cattle, horses, and many birds (data not shown). Mammals diversify VEGF-A by mRNA splicing. Teleost fish do the same but additionally have diversified VEGF-A by gene duplication. Zebrafish Vegfaa and Vegfab are both indispensable [101]. Vegfaa and Vegfbb isoforms of comparable length differ significantly in their charge, but it is unknown whether this translates into a differential interaction with the ECM.

Similarly to VEGF-A, VEGF-C might form gradients by the interaction of its SHD domain with the ECM [89]. Such morphogenetic gradients might be crucial for developmental lymphangiogenesis but also for developmental angiogenesis and vasculogenesis [51, 102,103], possibly explaining the strong purifying selection of VEGF-C in its VHD and SHD. Also, all other VEGFs, except for PlGF, showed strong conservation in the VHD (Supplementary Figure S5). For the VEGF-A isoforms, the sequence diversity outside the VHD increased with the length of the isoform (Supplementary Figure S3), perhaps facilitated by the existence of many isoforms.

### Structural differences between protostome and deuterostome PDGFs/VEGFs

PDGF/VEGFs form a family within the superfamily of cystine knot growth factors. Their hallmark is a characteristically spaced pattern of eight cysteine residues, consisting of the 6-cysteine pattern of the cystine knot signature expanded by two cysteines responsible for the covalent dimer formation of PDGFs/VEGFs [104]. The 8-cysteine pattern is broken with respect to the intermolecular disulfide bond-forming cysteine by only one vertebrate member of the PDGF/VEGF family, PDGF-C. However, in protostomes, missing intermolecular disulfide bonds are the rule rather than the exception (Figure 2). While disulfide bridges increase thermostability, low ambient water temperatures are typical for many freshwater and marine species, for which covalent dimer formation via disulfide bonds might not have any advantage over noncovalent dimer formation, while disulfide bond formation comes at a cost [105, 106]. Even at 37°C, the cystine bridge is not strictly necessary for dimer formation: VEGF-C also forms noncovalent dimers [19], and stable VEGF-A can also be produced after the mutation of the intermolecular cystine bridge-forming cysteines [107].

### Conserved when present, not needed when absent

Differently from VEGF-A and VEGF-C, which are pervasively maintained within the vertebrate lineage, PlGF and VEGF-B are absent from major vertebrate classes. PlGF appears to be absent in amphibians, and VEGF-B is completely missing from birds and crocodiles. The gene duplication that led to the establishment of the PlGF and VEGF-B genes presumably happened shortly before the cartilaginous fishes branched off. Consequently, e.g. shark VEGF-B is much more similar to VEGF-A compared to VEGF-B of land animals (Supplementary Figure S6).

The absence of PlGF in amphibians and VEGF-B in birds and crocodiles is due to secondary gene loss events, which were apparently – very similar to knockout experiments of the same genes in mice [12–14] – well tolerated. While VEGF-B has been proposed to play a role in the regulation of endothelial fatty acid uptake [108] and vascularization and tissue perfusion via indirect activation of VEGFR-2 [109], its precise role remains controversial [110, 111]. Its evolutionary loss might have been a net benefit for birds, perhaps even instrumental to enabling the high metabolic turnover needed for flight [112]. In any case, our understanding of PlGF, VEGF-B, and VEGF-D, all having been conserved for 500 MYA despite their apparent present-day redundancy in mice, leaves ample room for future insights.

While very common in plants, polyploidy is rare among animals. Among vertebrates, it is tolerated best by fish and amphibians [113]. This tolerance is also seen at the gene level. We frequently found individual *pdgf/vegf* gene duplications in fish but not in higher vertebrates. Holostei fish, a sister clade of the teleost fish, show, for example, a duplicated *vegfc* gene. Whether the duplicated *vegfc* genes have been maintained in Holostei from one of the prior whole genome duplications or whether they resulted from a limited gene duplication event early in the Holostei lineage is unknown and perhaps unknowable since the chromosomal context has likely been already lost. Surprisingly, both *vegfc* genes continue to be strongly conserved in Holostei. The conservation is strongest in the receptor binding domain but can also be seen in the SHD (Figure 6B). Interestingly, only two of the conserved residues of the PDGF/VEGF signature were under strong purifying selection, and only three out of the nine residues under strong purifying selection were cysteines, arguing that a better, perhaps more sensitive search pattern for the detection of PDGF/VEGF proteins could be developed by taking conserved non-cysteine residues into consideration. Contrasting this strong conservation is the variability of the immediately N-terminally adjacent region, which is presumably instrumental in the activation of the inactive pro-VEGF-C into the mature VEGF-C by proteolysis [114] (see also https://elifesciences.org/articles/44478/figures#fig2s2). Another gene duplication example is the loach *Triplophysa rosa* (Figure 6D), which features three *vegfa* and two *pgf* genes, both of which could be de-novo duplicated or maintained from one of the previous WGDs.

Evolutionary recent whole genome duplications have been reported for catostomid fishes such as the Chinese Sucker (*Myxocyprinus asiaticus*) [115, 116] and the common carp (*Cyprinus carpio*) [117, 118]. It would be interesting to analyze the pseudogenization pattern for PDGF/VEGF genes in these two species.

### Fish as model organisms

Teleost fish, which comprise most of the extant fish species, have undergone a lineage-specific whole genome duplication 350 MYA [52, 119], which resulted in presumably ten active *vegf* and eight active *pdgf* genes immediately after the duplication. In the teleost zebrafish, at least 12 of the duplicated *pdgf/vegf* genes remain functional until today, according to our analysis (*pdgfaa/ab, pdgfba/bb, pdgfc, pdgfd, vegfaa/ab, pgfb, vegfba, vegfc, vegfd*). While the Ensemble genome database also lists *<u>vegfbb</u>* as an active zebrafish gene, we did not find any mRNA transcript matching it in 21 fish species, including zebrafish. Similarly, we could not find any *pgfa* transcripts, despite being programmatically identified as an active gene by the Zebrafish Genome Reference Consortium (*<u>pgfa</u>*). It is unknown whether the genes represent pseudogenes or they were not expressed in the tissues that were used for mRNA extraction.

Different teleost lineages underwent distinct gene elimination patterns; this is the most simple explanation for the occurrence of, e.g., two *vegfc* genes in the European eel (order *Anguilliformes*) or the butterflyfish (order *Osteoglossiformes*). Salmonids, on the other hand, have undergone one additional full genome duplication approximately 88 MYA [54]. While this might have resulted in theoretically 36 different active *pdgf/vegf* genes immediately after the duplication, not all of these are active today. For the salmonid *Salmo trutta* (Brown trout), 26 of these genes are identified by the Ensemble analysis pipeline as active (https://www.ensembl.org/Salmo_trutta/Location/Genome?ftype=Domain;id=IPR000072), and for 21 of them, we found mRNA transcripts. However, if mRNA and protein data are absent, it is not always possible to reliably distinguish functional from pseudogenes [120].

The partial loss of gene function within the teleost lineage remains challenging for experimental fish – and specifically zebrafish – research. Multiple functions that were prior to the genome duplication executed by a single protein (e.g., VEGF-A) might be executed by two (zebrafish) or even more (Salmonidae, Catostomidae, *Cyprinus carpio*) ohnologs. There is multiple evidence that the two zebrafish ohnologs *vegfaa* and *vegfab* have diversified in terms of mRNA splicing and, consequently, their angiogenic properties [101, 121, 122]. Similarly, the ohnologs *vegfc* and *vegfd* have diversified differently in fishes compared to terrestrial animals in tissue distribution and receptor interaction [123, 124]. While this makes fish models at times more tedious and difficult to interpret compared to mouse models [125], it is at the same time a unique opportunity to discover and understand a morphological and physiological diversity unknown in mammals [51, 126].

## Materials & Methods

### Comprehensive database scan

BLAST searches were executed for a set of 13 reference proteins (human PDGF-A/-B/-C/-D, PlGF-3, VEGF-A_121/165/206_, VEGF-B_167/186_, VEGF-C/-D, and vammin-1) against the non-redundant NCBI protein database (corresponding to RefSeq Release 94). According to the Hit_def of each result, a hit was programmatically categorized based on the Hit_def field in the blast result as a *synonymous hit* (e.g., when the VEGF-D search results in a hit annotated with “VEGF-D” or “FIGF”), a *related hit* (e.g., when a VEGF-D search results in a hit annotated with “VEGF-B” or “PDGF”) or an *undefined hit* (e.g., when a search for VEGF-D results in a hit annotated with “hypothetical protein” or similar). To categorize undefined hits, secondary BLASTS were initiated with the sequences for undefined hits, and if more than 50% of the secondary hits agreed in their annotation on a specific PDGF/VEGF, this information was used to categorize the primary hit. The 50% threshold had been empirically determined to be conservative, i.e., never resulting in false negative categorizations with a known set of VEGFs. Undefined hits in secondary BLASTS results are assigned to specific PDGFs/VEGFs in the same fashion but using computationally more expensive RPS BLAST instead of protein blast. The flow chart of the analysis is shown in Supplementary Figure S1.

The full table of BLAST results (Supplementary Table 1) was assembled programmatically from the data generated as described above, and the number of distinct animal species for each clade was obtained from the NCBI taxonomy database. “Fully sequenced” genomes were loosely defined as those that had registered a BioProject with NCBI with the data type “Genome sequencing” or “Genome sequencing and assembly” in the group “animals” and for which results had been published (<u>1049 species</u> at the time of this writing). The number of protein sequences published for a specific clade was the number of sequences in the corresponding taxon-specific subsection of the NCBI protein database. For all clades with less than 200 unique VEGF homologs, all primary BLAST hits were manually checked for false positives (i.e., when the human-curated protein description specified a named non-PDGF/VEGF protein in the sequence description). The formula for background coloring of Figure 1 and Supplementary Table 1 according to the heuristic reliability due to biased sampling was: 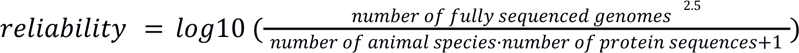

### Alignment, phylogenetic tree building, and conservation analysis

#### Alignment of cnidarian with human PDGFs/VEGFs (Figure 2)

The mcoffee mode of T-coffee 12.00 was used to align a representative subset of 10 cnidarian PDGF/VEGF-like sequences from all six species in which PDGF/VEGF-like sequences were identified, all seven human PDGF/VEGF orthologs, *C. elegans* PVF-1, and *Drosophila* PVF2. The alignment was trimmed down to 130 amino acid residues of the consensus sequence (corresponding to the VHD) starting from the first of the eight conserved cysteines of the PDGF/VEGF signature (until Proline-132 from VEGF-A). During the alignment, all conserved cysteine residues were anchored according to the alignment by Heino et al. [35].

#### Assessing the relationships between vertebrate and invertebrate PDGFs/VEGFs (Figure 3)

The alignment from Figure 2 above was expanded by including all four PDGFs/VEGFs identified in *Hydra vulgaris*, all three *Drosophila* PVFs, all human PDGF/VEGF paralogs, and the PDGF/VEGF-like molecule identified in the parasite *Brugia malayi*. The TGF-β homolog of *C. elegans* UNC129 was used as an outgroup. To capture information about the ancillary domains/propeptides of the proteins, the sequences included in the alignment were expanded amino-terminally by 20 amino acids and C-terminally by 30 amino acids beyond the conserved eight cysteine residues of the PDGF/VEGF signature. The mcoffee-provided alignment was trimmed to include 179 amino acid positions corresponding to the first amino acid of the major mature form of human VEGF-C [19] until the last cysteine of the first repeat of the BR3P motif (C-X_10_-C-X-C-X_(1,3)_-C) [15]. Tree building was performed with PhyML 3.0 [127], combining both the original PhyML algorithm and subtree pruning and regrafting for tree topology search. To estimate the reliability of branches, 1000 bootstrapping replicates were performed. The tree was visualized with FigTree version 1.4.4, exported to an SVG file, and visually enhanced using Inkscape (https://inkscape.org).

#### Holostei VEGF-C analysis (Figure 6A-C)

The amino acid alignment and tree building for Holostei VEGF-Cs and VEGF-Ds were performed as described above using T-coffee and PhyML. As an outgroup sequence for tree rooting, we used PDGF-A from one of the Holostei species (*Lepisosteus oculatus*). The corresponding mRNA alignment was generated with PAL2NAL. Treefile and alignment were used by HyPhy version 2.5.1 [128] to test for pervasive site-level selection (SLAC). The graphs were generated from the json files using Gnumeric. The alignment of the VHD of representative VEGF-C sequences below the detailed conservation view was generated from an aligned Fasta file with SnapGene Viewer 6.1.1.

#### Tree building for Triplophysa rosa PDGFs/VEGFs (Figure 6D)

All 11 PDGF/VEGF sequences identified for the sucker *T. rosa* were aligned, and tree building was performed as described above. For this analysis, we did not include any outgroup resulting in an unrooted tree.

#### Analysis of viral VEGF homologs

A PSI-BLAST limited to the taxon Viridae (taxid:10239) was run against the starting sequence AAD03735.1 (vascular endothelial growth factor homolog VEGF-E from Orf virus) until no new sequences were found above the 0.005 threshold. The Fasta descriptions were adjusted to include virus names and host species. For each host species, the VEGF-A_165_, PlGF-1, and VEGF-B186 orthologous protein sequences were retrieved (if available) and included in the alignment and tree building, which was performed as described above. Only three out of the eight fish VEGF-B sequences were available. Therefore, we included both zebrafish VEGF-B sequences in the analysis. Similarly, no VEGF sequences were available for *Halichoerus grypus* (grey seal). These were replaced with sequences from the closest species for which VEGF sequences were available (*Zalophus californianus*, California sea lion). The protein sequence alignment was performed with T-coffee, the tree building with PhyML, and the visualization of the tree with the ETE Toolkit 3.0. The v-sis sequences from the Simian sarcoma virus were used as an outgroup to root the tree. The workflow and all sequences used for the analysis are available from GitHub as a python script (https://github.com/mjeltsch/VEGFE). Tree topology was used to infer the likely origin(s) of viral VEGFs.

#### Analysis of VEGF-F origin

A tree was constructed with T-coffee and PhyML using 15 snake venom VEGFs and the hemangiogenic VEGFs from four bird species, eight mammals, eleven reptiles, and two amphibians, for which a complete set of proteins was available (excluding VEGF-B in birds and crocodiles and PlGF in amphibians). The tree was visualized with iTOL v5 [129] and enhanced using Inkscape.

#### Comparison of conservation levels between individual VEGFs

To compare the degree of conservation and to identify possible positive selection, codon analysis (synonymous versus nonsynonymous changes) was deployed. Reference protein and transcript sequences for available ortholog sets were downloaded from NCBI (https://www.ncbi.nlm.nih.gov/gene/XXXX/ortholog, XXXX = 7422, 7423, 7424, 2277, 5228 for VEGF-A, -B, -C, -D and PlGF, respectively). Obviously bogus or truncated transcript predictions (lacking essential exons of the PDGF/VEGF homology domain or being of low quality) were eliminated manually from the set. Several proteins/transcripts had to be replaced manually to ensure that only transcripts of the same isoform were compared since not all species feature the same set of isoforms (e.g., for VEGF-C: Sus scrofa, for VEGF-A: Mus musculus). Species were only included if they featured the full set of five mammalian VEGFs in the database (VEGFA, PlGF, VEGF-B/-C/-D). The full list (sets of 5 VEGF paralogs for 82 species) was reduced to comprise only the 50 most informative sequences using T-coffee [130]; however, all sequences in a set were maintained if only one sequence in a set had been classified as informative. The full set of protein and mRNA sequences is available in Fasta format as Supplementary data. The protein sequences of each VEGF ortholog were aligned using T-coffee’s mcoffee mode [131]. The final alignments were trimmed manually (keeping the sequence from the first to the last cysteine of the PDGF/VEGF cysteine signature plus 15 amino acid residues N-terminally and 5 amino acid residues C-terminally. To prepare the protein alignment for the analysis of pervasive purifying/adaptive evolution, the corresponding mRNA alignment was obtained with PAL2NAL 14 [132]. A maximum-likelihood (ML) approach was used to infer nonsynonymous versus synonymous substitution rates on a per-site basis for the alignment [133], using SLAC (Single Likelihood Ancestor Counting) from the HyPhy 2.5.1 software package. The graph was generated with the Python library Seaborn 0.10 and enhanced via Inkscape with a molecular ribbon model of VEGF-C generated by Pymol 2.3.0 based on the PDB structure 2X1X.

### Cladograms of evolutionary PDGF/VEGF history

A tree file for the animal kingdom was downloaded from the Open tree of Life (https://tree.opentreeoflife.org), which maintains a consensus tree obtained by semi-automated synthesis of many individual studies [44] (https://tree.opentreeoflife.org/curator). This treefile was used programmatically by the ETE Toolkit to generate Figure 1 and Supplementary Table 1. To generate Figure 4, a cladogram was generated from the same file, excluding the protostome branch with FigTree version 1.4.4. The PDF-exported tree was enhanced using Inkscape. When public domain animal silhouettes were available, they were obtained from PhyloPic (http://phylopic.org); otherwise they were drawn by the authors.

### Determination of absence of VEGF-B genes in birds

Because we had not come across any avian VEGF-B sequence, we started by blasting the Entrez protein database (ref) with a reptilian VEGF-B protein sequence (Chinese soft shell turtle VEGF-B, Uniprot K7FWR8). The sequence ids of all hits were re-written to include the animal class and common name of the organism to facilitate the identification of potential avian VEGF-B sequences. This set of 4976 sequences was subjected to multiple sequence alignment (MSA) using mcoffee. Gblocks was used with the following in order to manually curate the alignment [134]. After converting the output with Dendroscope 3.8.4 to Newick format, the graphical representation was generated with iTOL [135], visually refined using Inkscape, and inspected for avian VEGF-B sequences. In order not to miss distantly related sequences, we repeated the above analysis but used three rounds of a PSI-BLAST, based on all available VEGF-B sequences and all avian VEGF-A and PlGF sequences).

### Fish mRNA analysis

The PhyloFish mRNA database (http://phylofish.sigenae.org) was queried using tblastn with all *Danio rerio* PDGF/VEGF protein sequences, resulting in 1547 unique transcripts. mRNA sequences were downloaded for all transcripts. 405 of these transcripts were identified as PDGF/VEGF family members with a relaxed PDGF signature (using the regular expression P.?C.{2,8}C.?G.?C). Individual phylogenetic trees were built for the PDGF/VEGF transcriptome of each species using *Danio rerio* PDGFs/VEGFs as reference sequences. Unannotated mRNA sequences were classified manually based on the nearest reference sequence neighbor on the tree. The total number of unique mRNA transcripts for each species and the number of unique transcript contigs were tabulated using Gnumeric version 1.12.46 (Supplementary Table 3). The numbers were normalized to zebrafish, for which 48158 unique mRNA transcripts had been obtained. The data was visualized using the Gnumeric Matrix plot function. The plot was visually enhanced in Inkscape with a cladogram based on [52]. For the extended mRNA analysis (Supplemental Figure S7), which compares mRNA levels of different organs within one fish species, expression levels were not normalized.

### Quantification of diversity

As the DIVAA software could not be retrieved anymore, we reimplemented the algorithm in Biophython based on the publication [69]. As an input to the DIVAA algorithm, VEGF-A isoforms were programmatically determined, but VEGF-B isoforms were manually assigned because fishes were found to have a single VEGF-B isoform which displays features of both mammalian isoforms simultaneously, having a basic stretch C-terminally to the VHD, followed by a hydrophobic region and a terminal, basic, cysteine-rich region. For other VEGFs, different isoforms were not separately analyzed.

## Declarations

## Acknowledgements

We thank Jeremy Pasquier et al. for access to the fish mRNA sequencing data. We also wish to acknowledge CSC – IT Center for Science, Finland, for the provisioning of computational resources.

## Availability of data and material

All of the data used for this study are available from the corresponding GitHub repositories. The original search results are online at: https://mjlab.fi/3Hm8PaB9Kee_usD1xll/animalia.svg.

## Code availability (software application or custom code)

Scripts and algorithms used in this study are available from the following GitHub repositories:

● https://github.com/mjeltsch/VEGFE
● https://github.com/mjeltsch/Holostei
● https://github.com/mjeltsch/VEGFphylo
● https://github.com/mjeltsch/cnidariaVEGFs
● https://github.com/mjeltsch/VEGFselect
● https://github.com/mjeltsch/Fish_mRNA
● https://github.com/mjeltsch/divaa

## Authors’ contributions

KR and MJ analyzed and interpreted the data, drafted, wrote, and edited the manuscript. MJ developed and performed the bioinformatics analyses, acquired funding, and supervised the study. HB performed the phylogenetic analyses. KR manually curated programmatically uncategorized data.

## Supplementary Materials

**Supplementary Figure S1.**
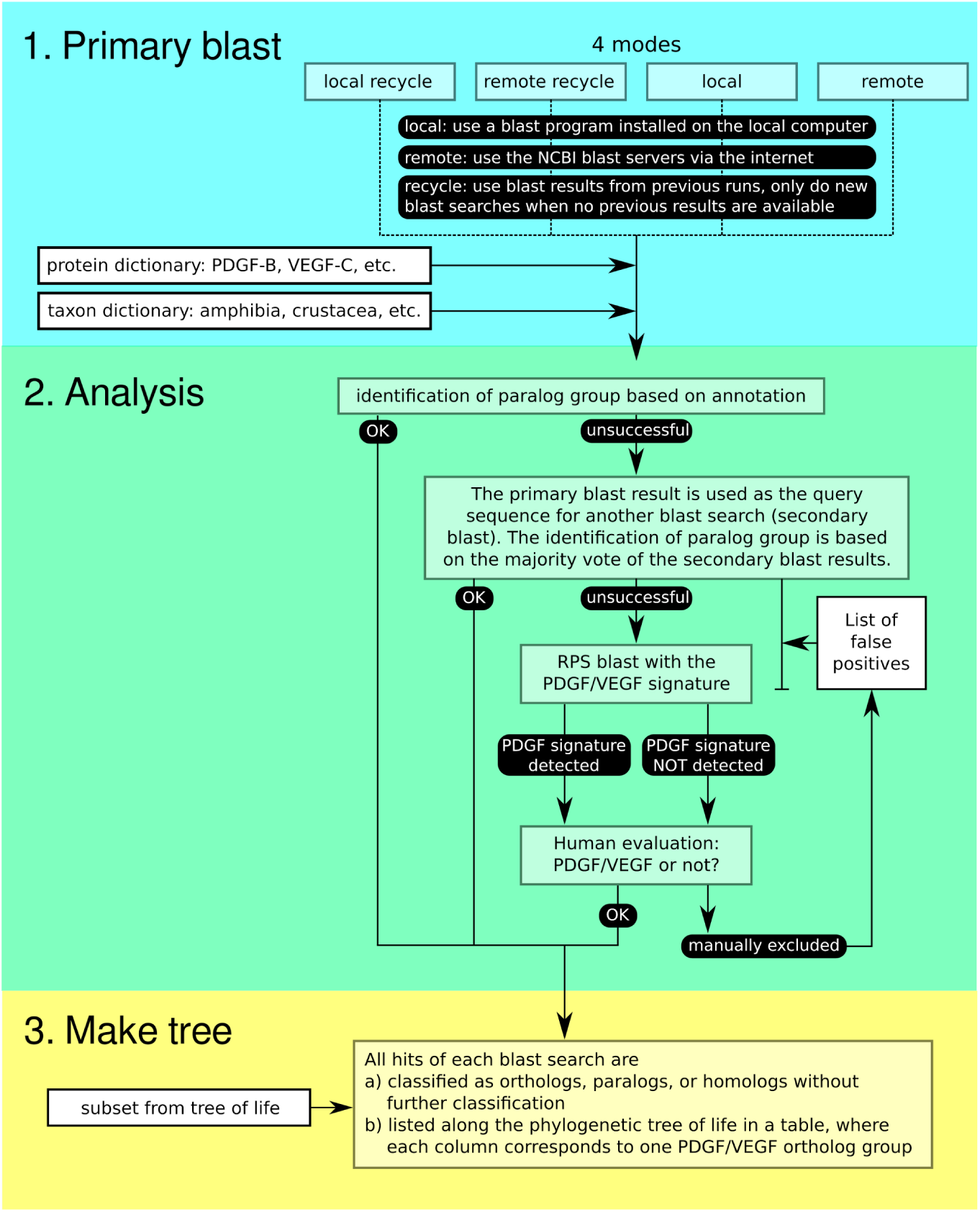
Bioinformatics workflow of the PDGF/VEGF search. The workflow resembles crowdsourcing by recursively analyzing the manual and programmatic descriptions of protein database entries most similar to the protein of interest. Only if this method fails, the protein is classified as a “generic” PDGF/VEGF homolog based on the detection of the PDGF signature (https://prosite.expasy.org/PDOC00222).

**Supplementary Figure S2.**
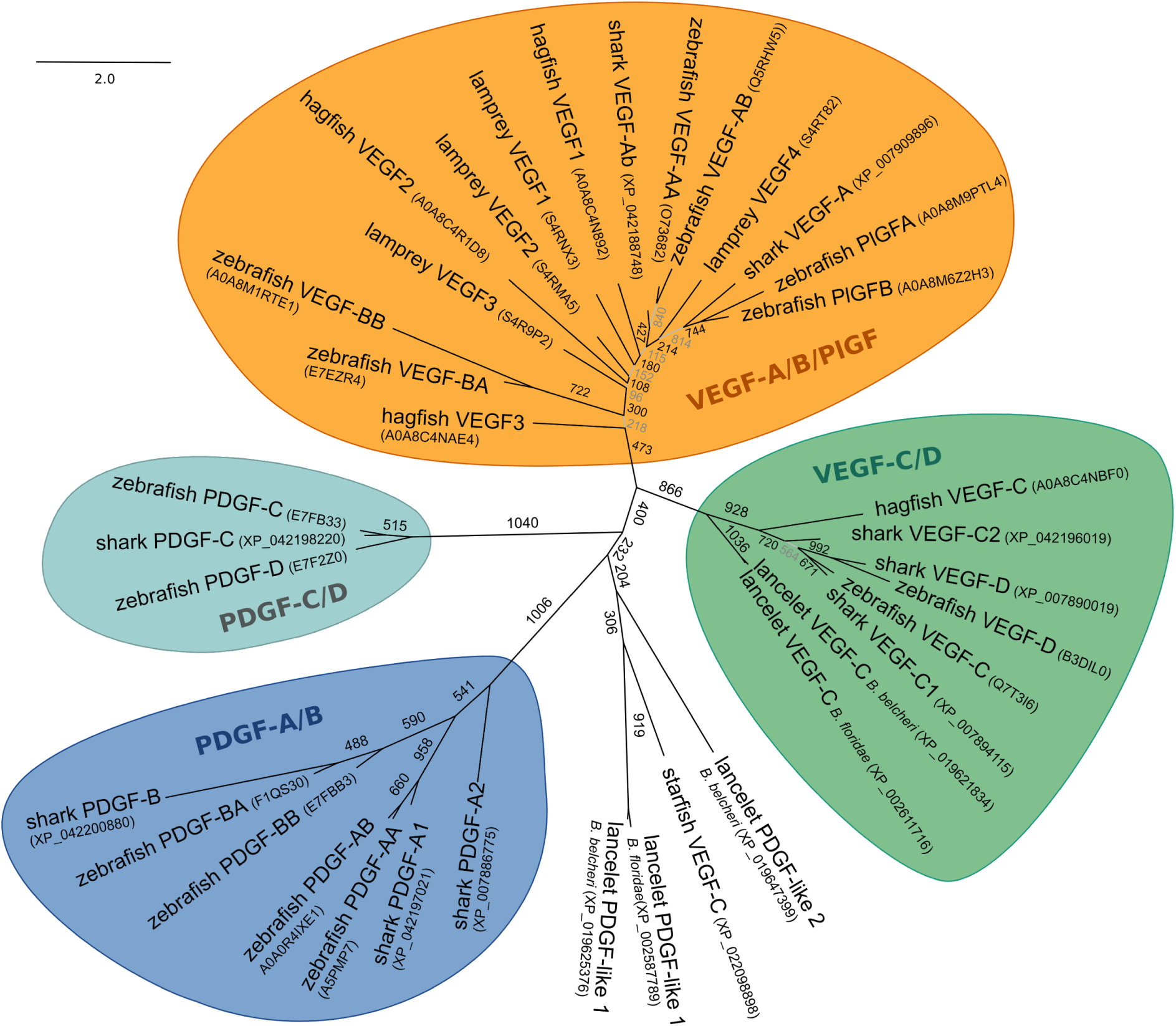
Relationship of early fish PDGFs/VEGFs. An unrooted tree of the relationships of PDGFs/VEGFs found in Florida and Chinese lancelets (*Branchiostoma floridae*, *Branchiostoma belcheri*), the inshore hagfish (*Eptatretus burgeri*), the sea lamprey (Petromyzon marinus), the elephant shark (*Callorhinchus milii*) and zebrafish (*Danio rerio*) shows that many of the early vertebrate PDGFS/VEGFs cannot be easily designated as orthologs of any modern VEGF or PDGF. Only with the appearance of bony fish, it becomes consistently possible to identify orthologs of these factors among the nine mammalian PDGFs/VEGFs. While there are clear VEGF-C-like proteins in jawless fish, most of their PDGF/VEGF-like molecules are difficult to classify. In this tree, such PDGFs/VEGFs are shown to be most closely related to VEGF-A/VEGF-B or PlGF, but bootstrap supports are generally weak. Even less clear are the PDGFs/VEGFs of Echinodermata and Cephalochordata, which appear equally distant to all the big groups (PDGFs, VEGF-C/D, VEGF-A/B/PlGF). Therefore, the assignment of PDGFs/VEGFs more distantly related to mammals than bony fish (such as shown in Figure 4) is hypothetical and relies mostly on the presence (VEGF-C-like) or absence (VEGF-A-like) of multiple BR3P repeats of BR3P motif.

**Supplementary Figure S3.**
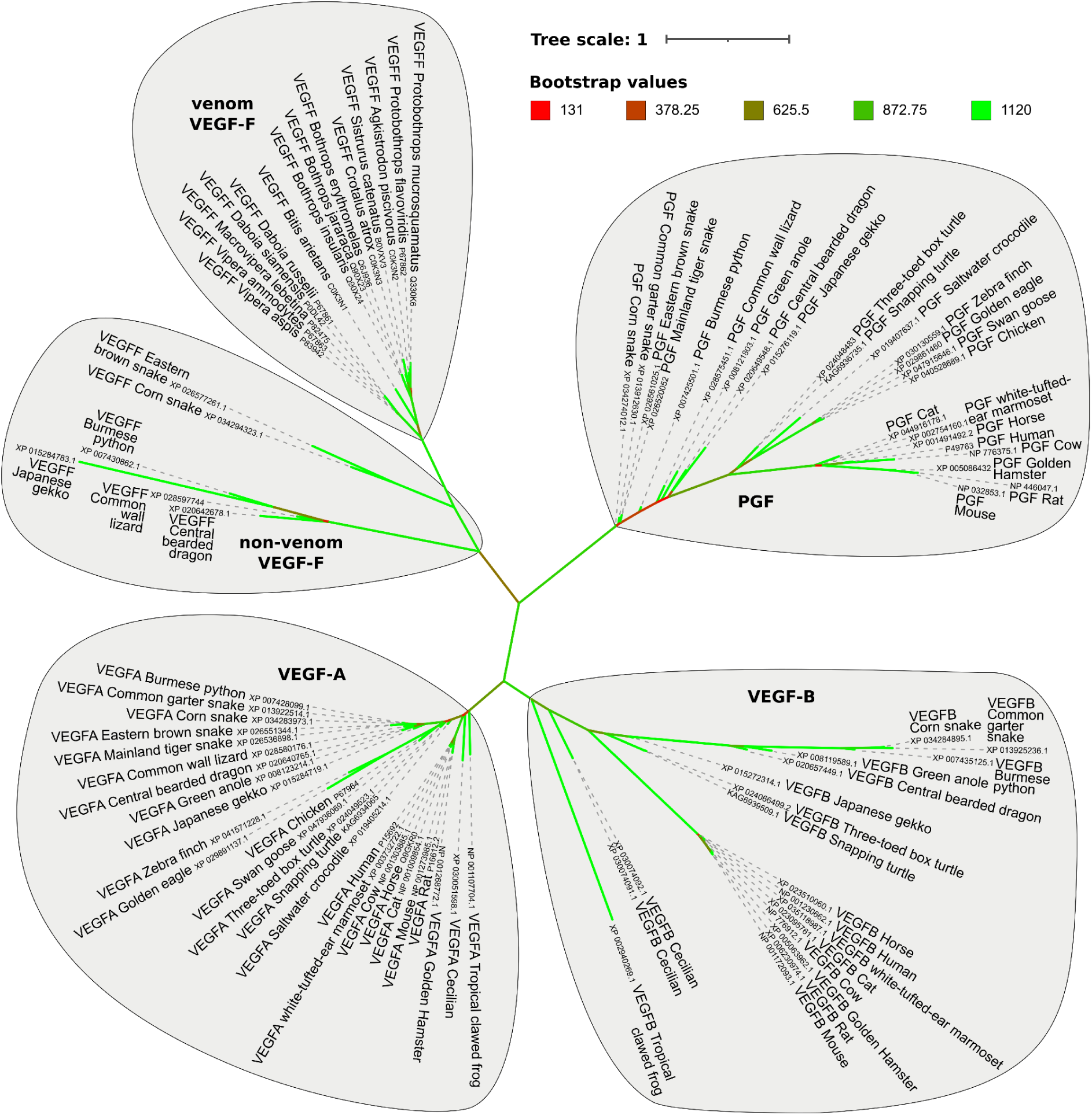
Phylogenetic tree of VEGF-Fs. An unrooted tree of the relationship of VEGF-F to the other hemangiogenic VEGFs (VEGF-A, VEGF-B, PlGF). VEGF-Fs segregate into multiple branches, supporting the two-origin hypothesis of venom (once in snakes, once in lizards, [78]). Assuming the prevailing two-origin hypothesis of venom, the independent emergence of VEGF-F as a venom component on multiple branches appears unlikely; even more so since several snake VEGF-Fs segregate with the non-venom (“lizard”) sub-branches. While venom VEGF-F expression is specific to the venom gland [78], it is not ubiquitous among venomous snakes and is found only in the viper clade (see also Supplementary Figure S14 in [78]). The most parsimonious explanation is a single origin of VEGF-F, predating the evolution of venom. Its original function is yet to be determined, but a later co-option of the VEGF-F gene in vipers established it as a bonafide venom component. However, in other branches of the tree (lizards and snakes outside the viper clade, such as the non-venomous Burmese python), the VEGF-F gene might serve its unknown original function, assumed a function as a venom component, or has disappeared. The most likely origin of the VEGF-F gene is a VEGF-A gene duplication. An origin via a PGF or VEGF-B gene duplication is less likely, but not impossible.

**Supplementary Figure S4.**
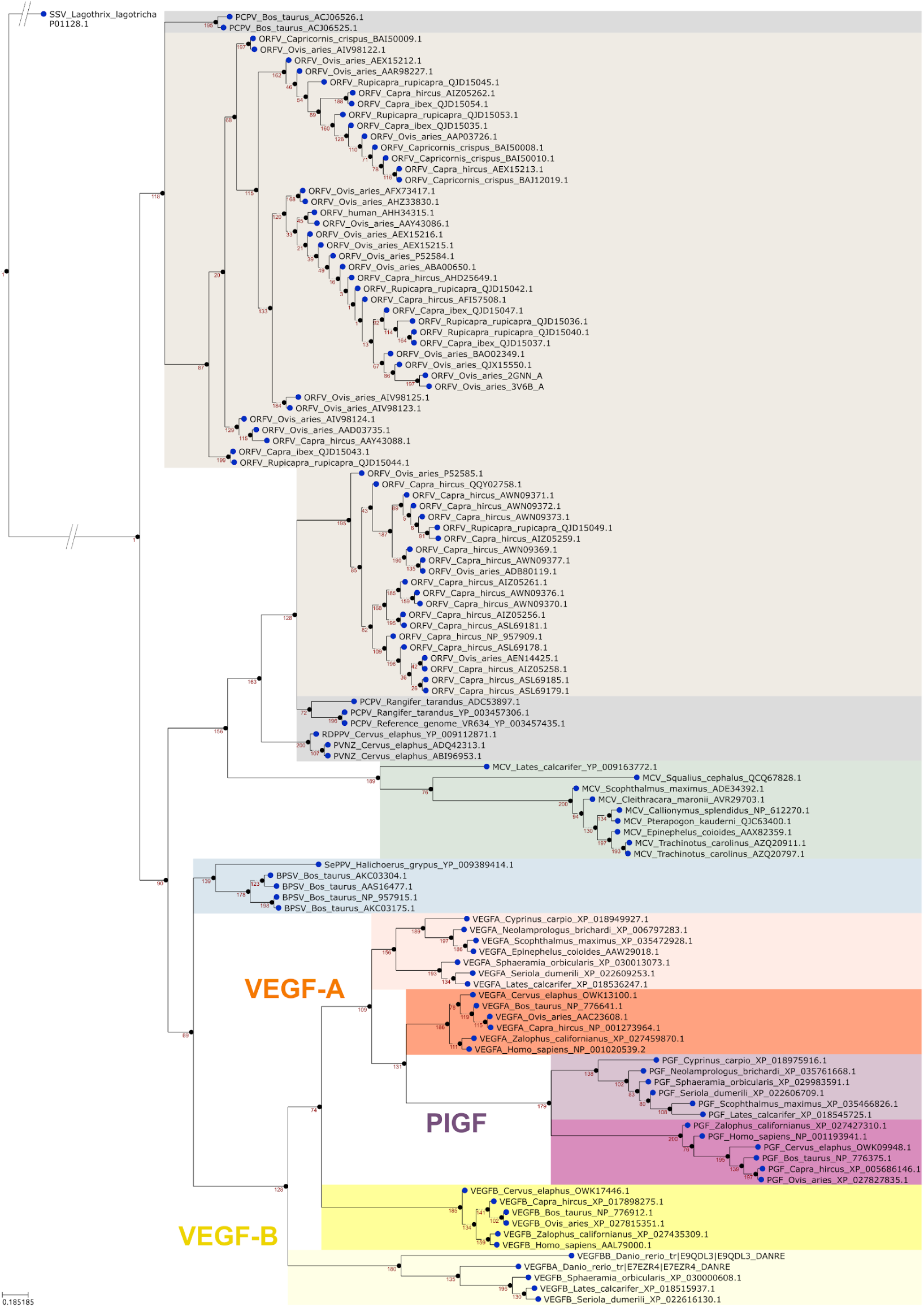
Uncollapsed phylogenetic tree of the VEGF-E tree ￼￼ in Figure 7. The most likely candidate for the VEGF-E origin is a single VEGF-A gene acquisition event from a vertebrate host.

**Supplementary Figure S5.**
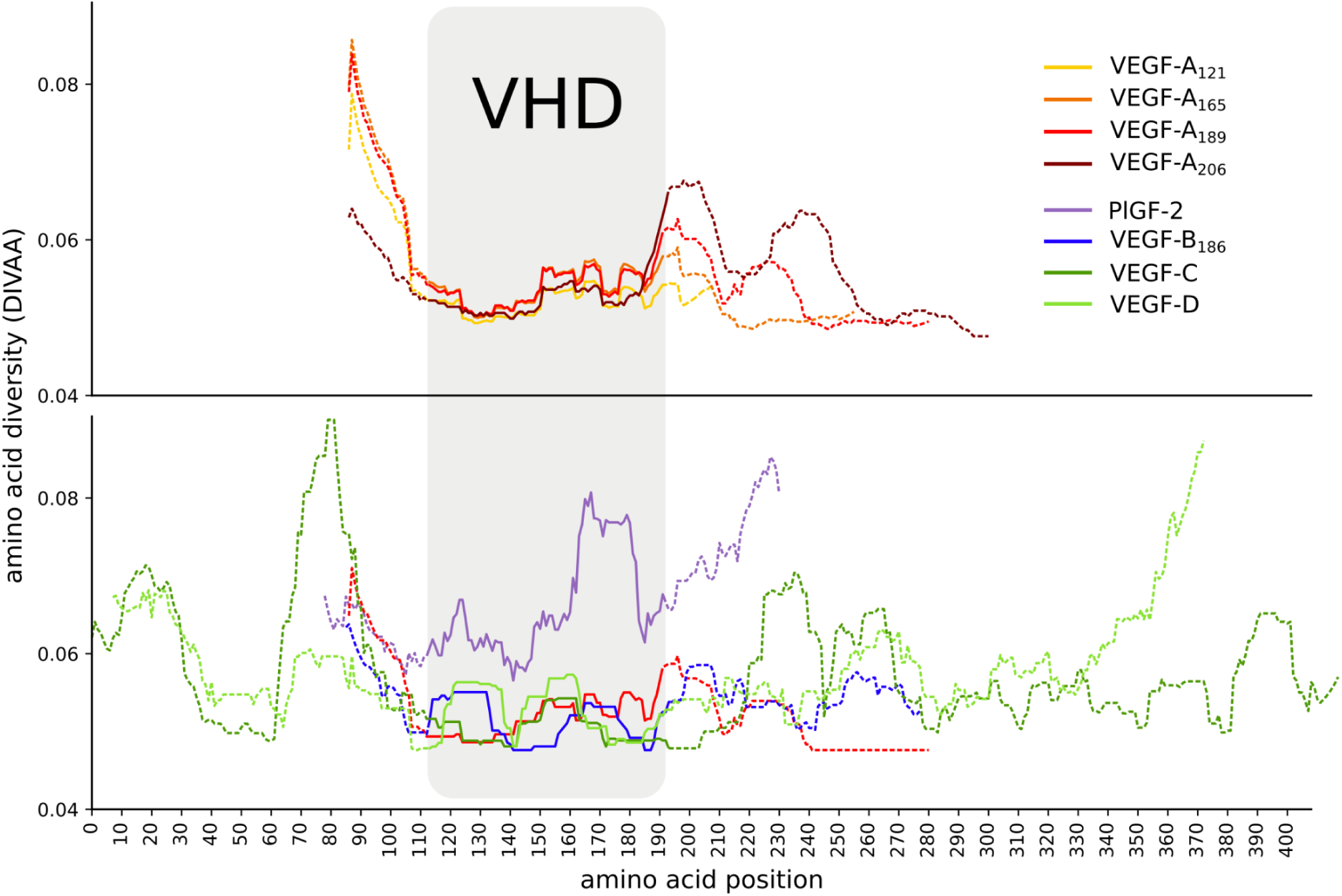
DIVAA analysis of VEGF-A isoforms (top panel) and the other mammalian VEGF family members (bottom panel). Although VEGF-A_165_ is the major isoform in humans, database entry frequencies indicate that VEGF-A_189_ might be the predominant VEGF-A isoform in many species. The diversity of all VEGFs is lowest in the VHD (i.e., the receptor binding domain) and increases both N- and C-terminally. From the VEGF family members, PlGF-2 shows the highest and VEGF-C the lowest level of diversity in the VHD.

**Supplementary Figure S6.**
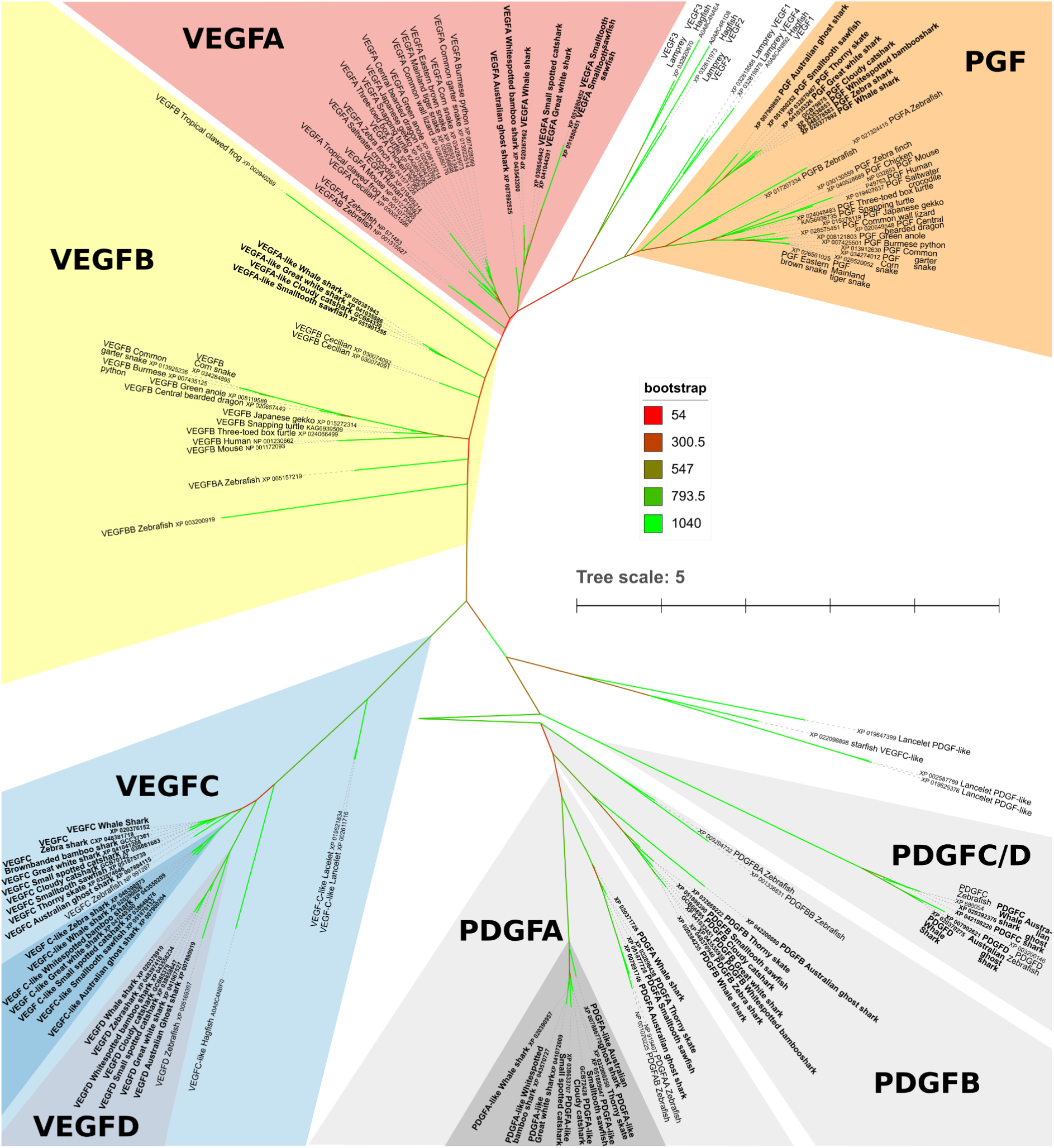
Sharks feature PlGFs and VEGF-B, and duplicated *vegfc* and *pdgfa* genes. Cartilaginous fish (chondrichthyes) arguably feature already orthologs to all nine mammalian PDGFs/VEGFs. However, since the separation of VEGF-A, VEGF-B and PlGF had been a fairly recent event when chondrichthyes separated from bony fishes, a clean assignment of individual genes into the VEGF-B versus the VEGF-A paralog group is ambiguous (these genes are labeled as VEGF-A-like in protein databases). Like zebrafish VEGF-B, these genes do not produce two transcripts with overlapping reading frames characteristic of mammalian VEGF-B. The C-termini of the translated proteins are highly positively charged similar to the human VEGF-B_167_ and VEGF-A_165_ isoforms. Chondrichthyes’ VEGF-A genes are often labeled as VEGF-Ab or VEGF-Ab-like, even though cartilaginous fish - different from zebrafish - did not undergo the teleost whole genome duplication. Interestingly, all sharks analyzed featured a duplicated VEGF-C and PDGF-A gene (“VEGFC-like” and “PDGFA-like”). PDGFs/VEGFs from cartilaginous fishes are shown in bold. For VEGF-C, VEGF-D, and PDGFs, the corresponding tetrapod proteins are not shown. With one exception, lancelet and cyclostomate proteins are not assigned to ortholog groups due to the low confidence in the branching. Confidence values were determined by bootstrap analysis, and are shown in a color scale from red (low confidence) to green (high confidence).

**Supplementary Figure S7.**
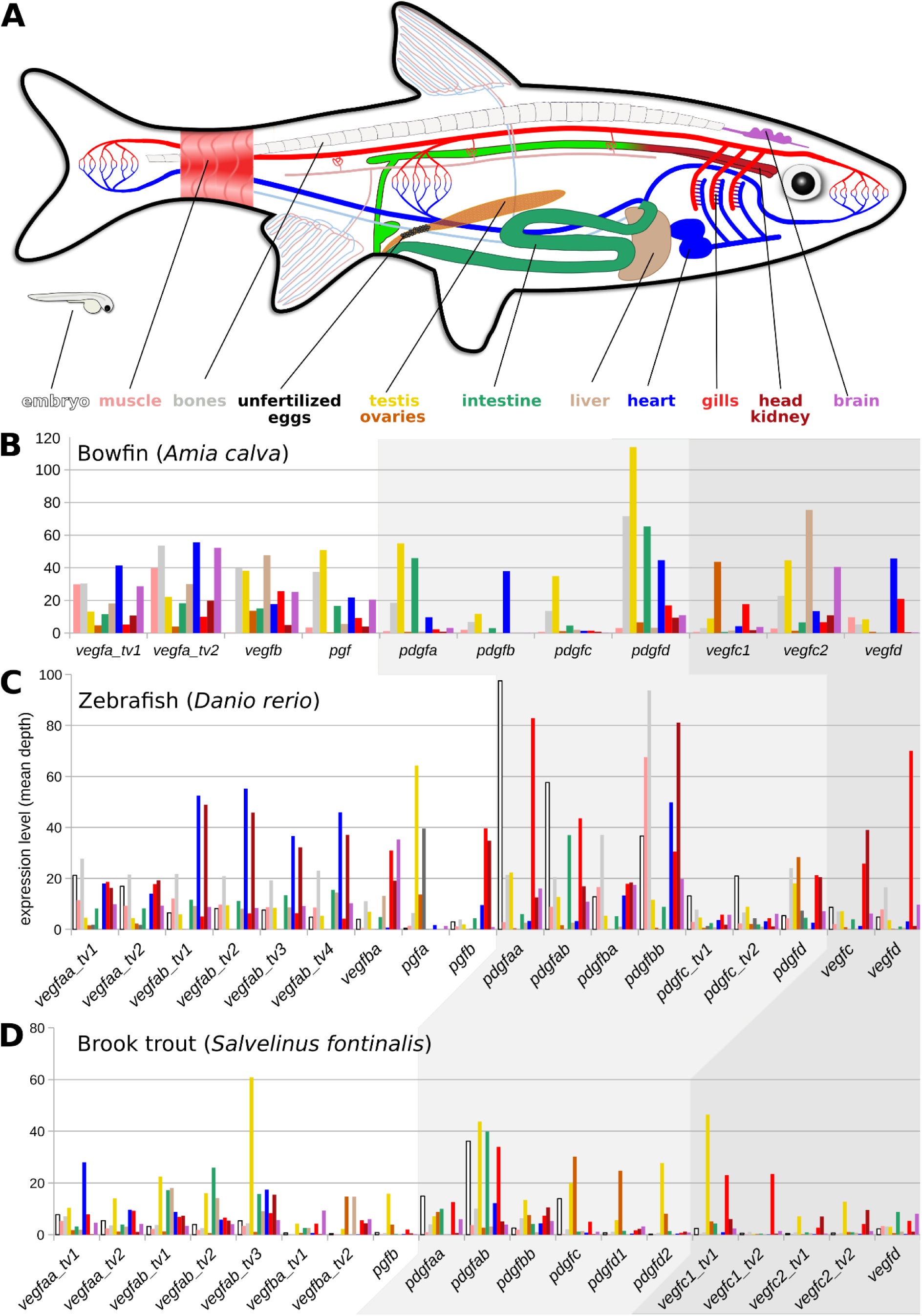
Expression of PDGFs and VEGFs in bowfin, zebrafish, and brook trout. The bowfin has undergone none, zebrafish one, and brook trout two whole genome duplications after the second common vertebrate genome duplication. The analyzed tissues are color-coded, as shown in (A). The _tv# extension designates different transcript variants (e.g., splice isoforms). Genes that have been duplicated independently from the teleost genome duplication have been indicated by adding a numeral after the gene name.

**Supplementary Table 1.** Complete table of all BLAST hits.

**Supplementary Table 2.** False-positive BLAST hits of PDGF/VEGF-like sequences from the phylum Porifera.

**Supplementary Table 3.** Fish PDGF/VEGF mRNA transcript contigs.

**Supplementary Table 4.** Complete list of all 24 fish mRNA data.

