## Supplementary figures and images for "Expansion and collapse of VEGF diversity in major clades of the animal kingdom"

### Supplementary Figure 1

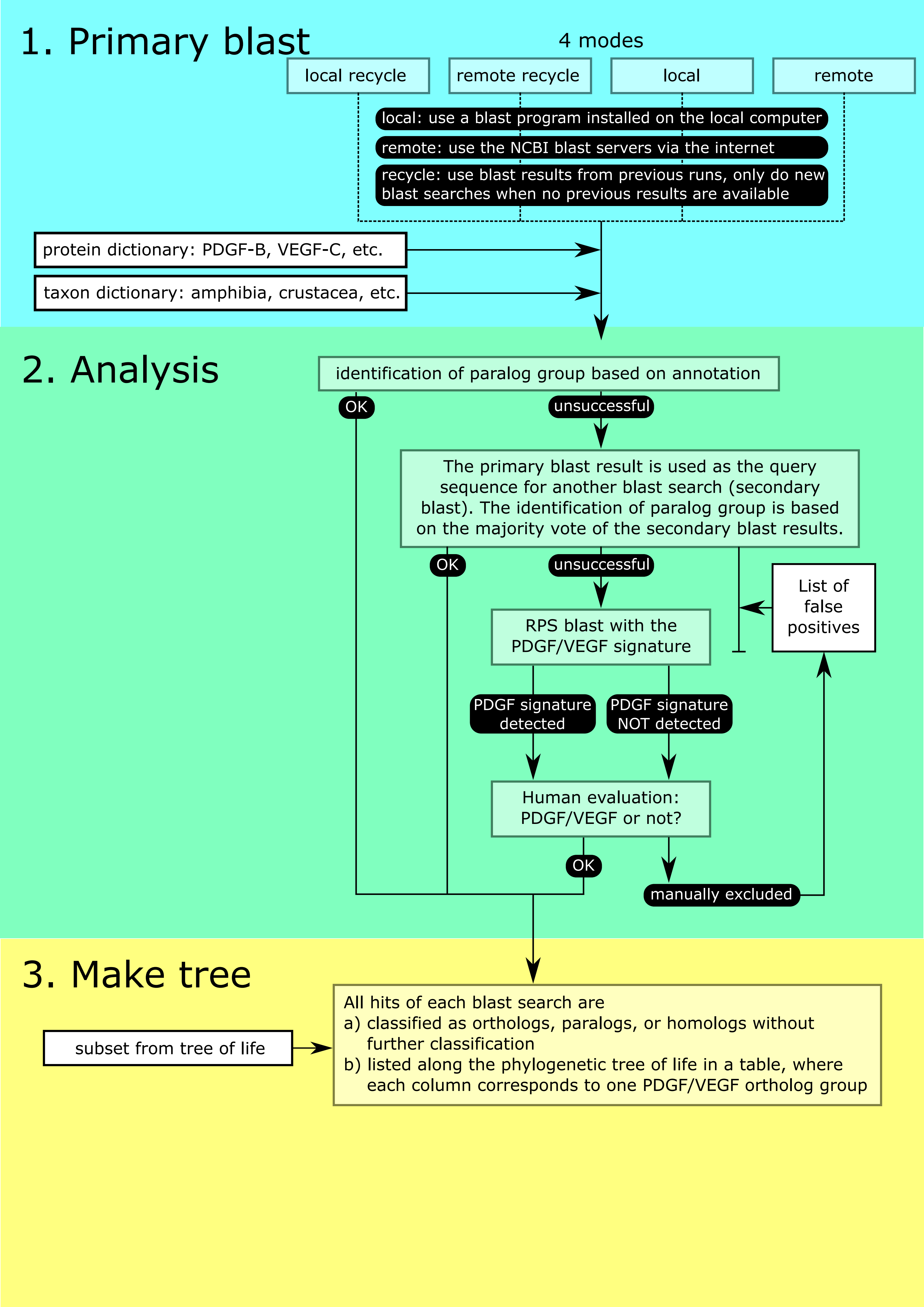

### Supplementary Figure 2

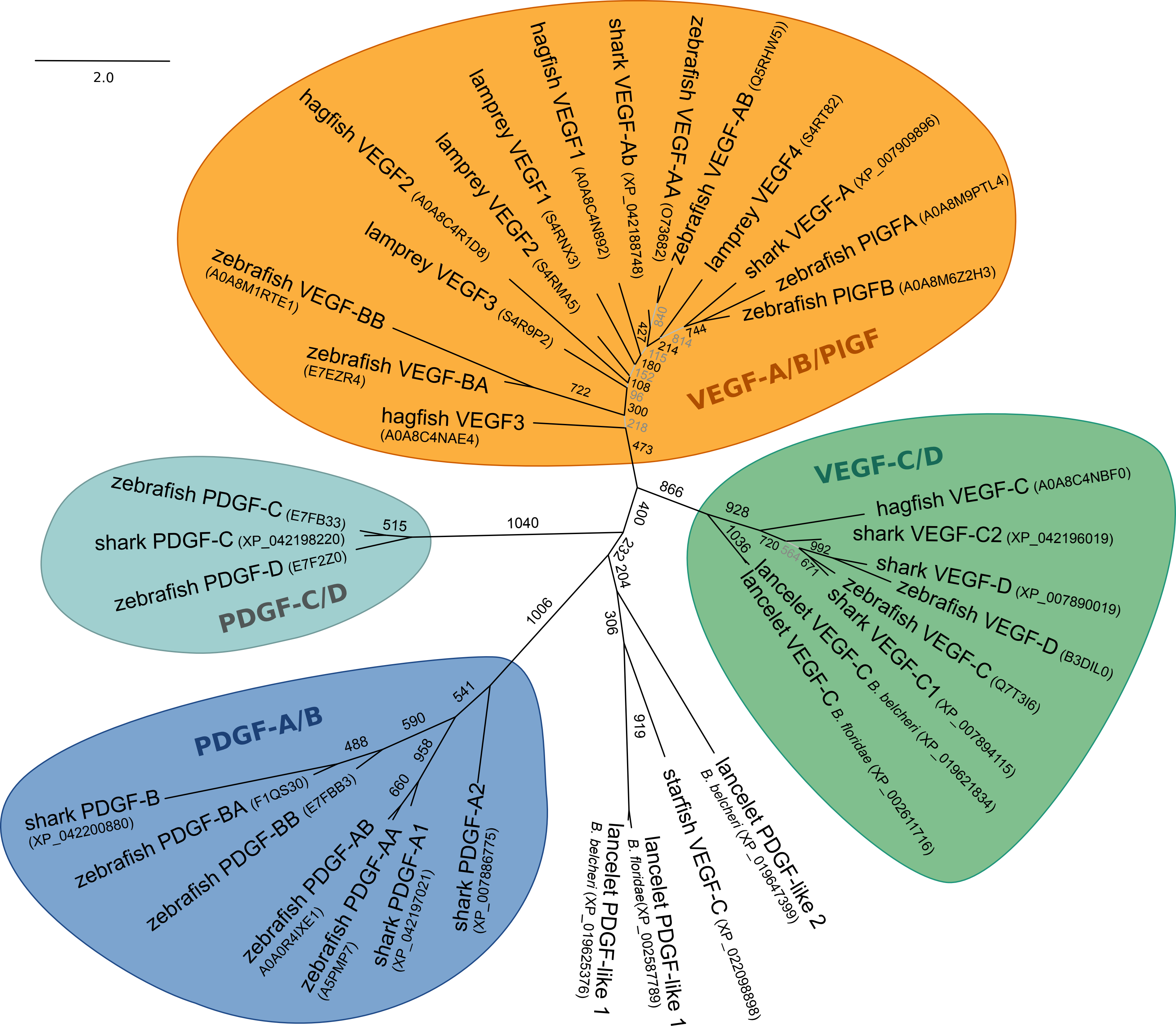

### Supplementary Figure 3

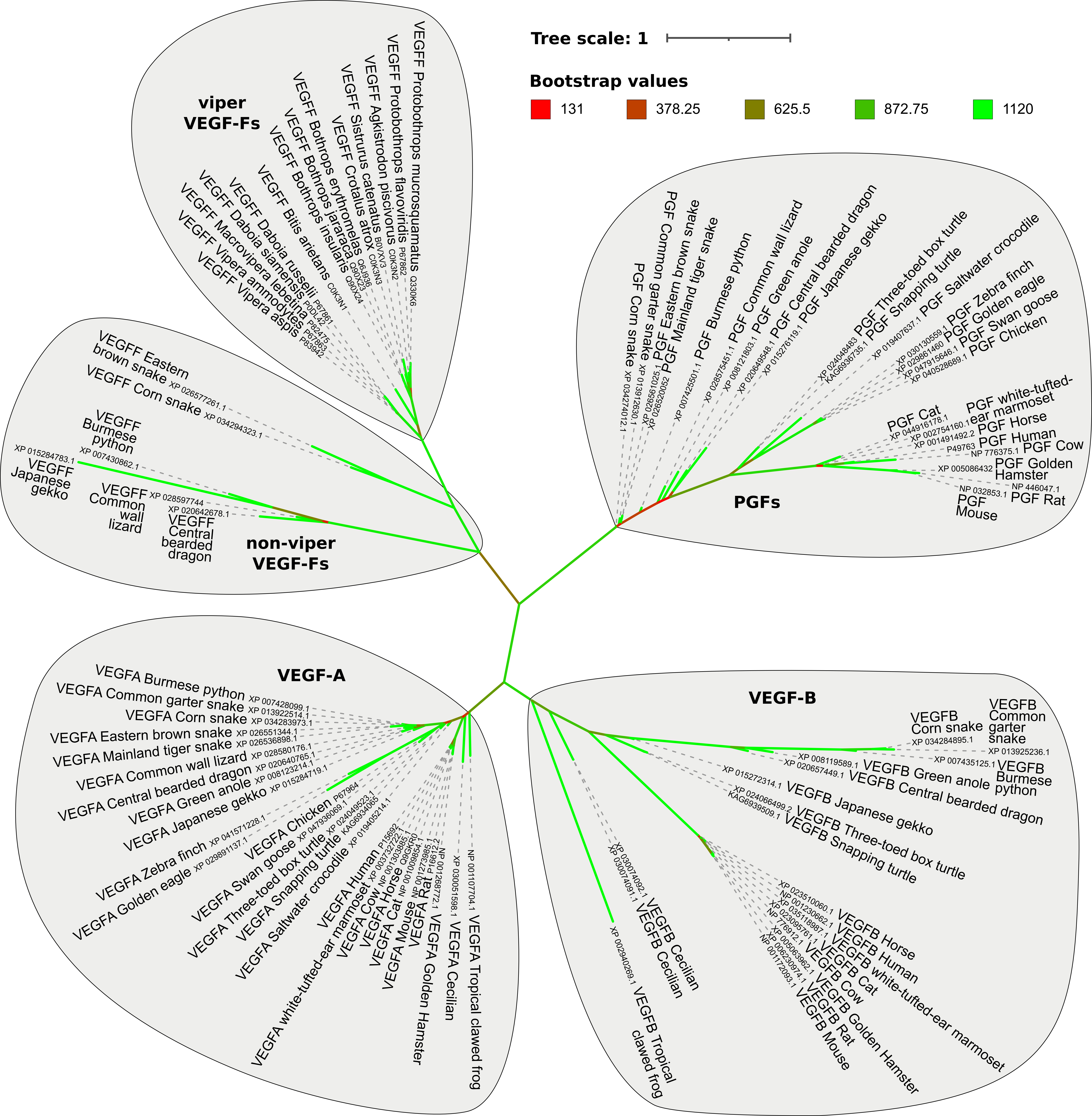

### Supplementary Figure 4

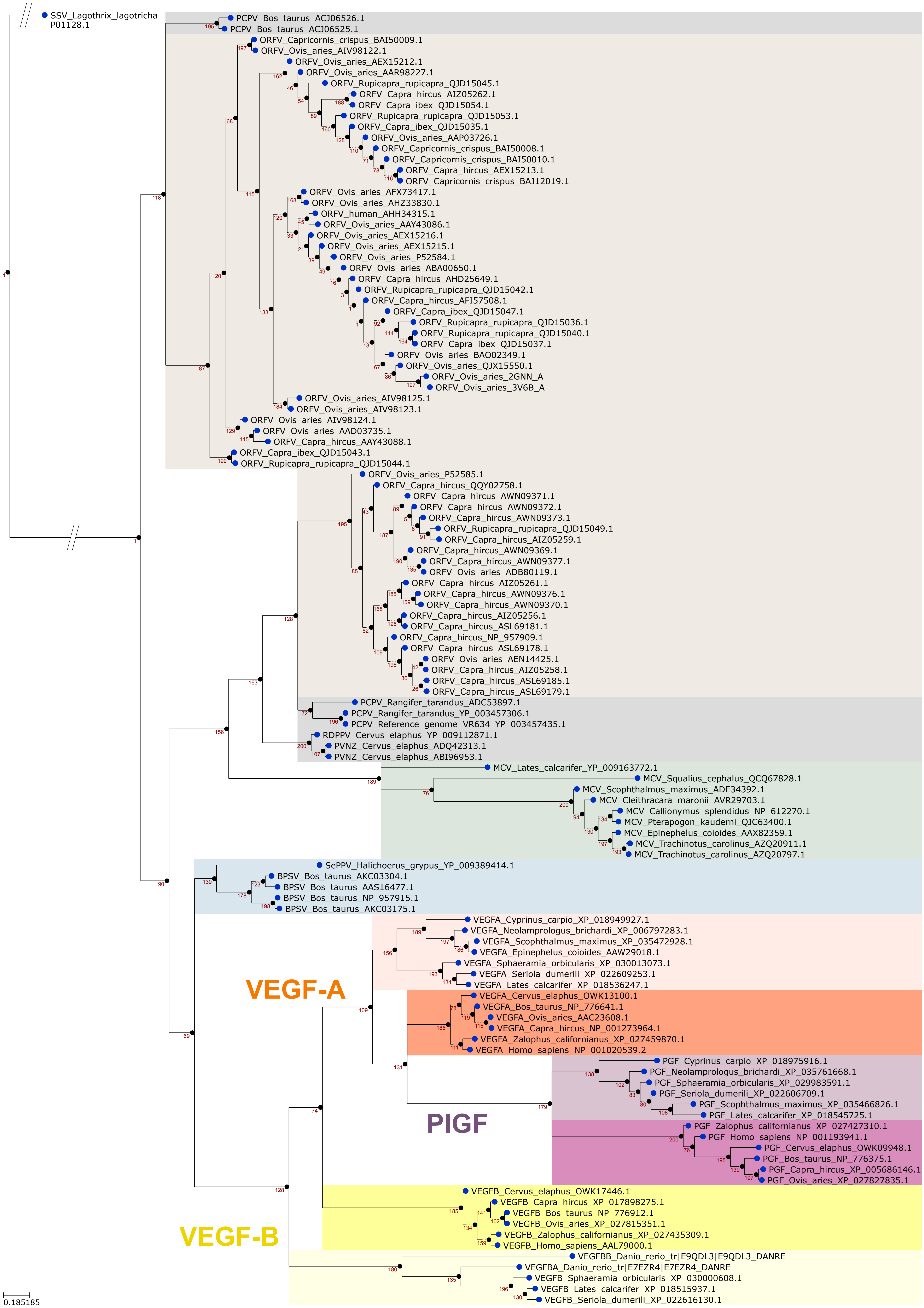

### Supplementary Figure 5

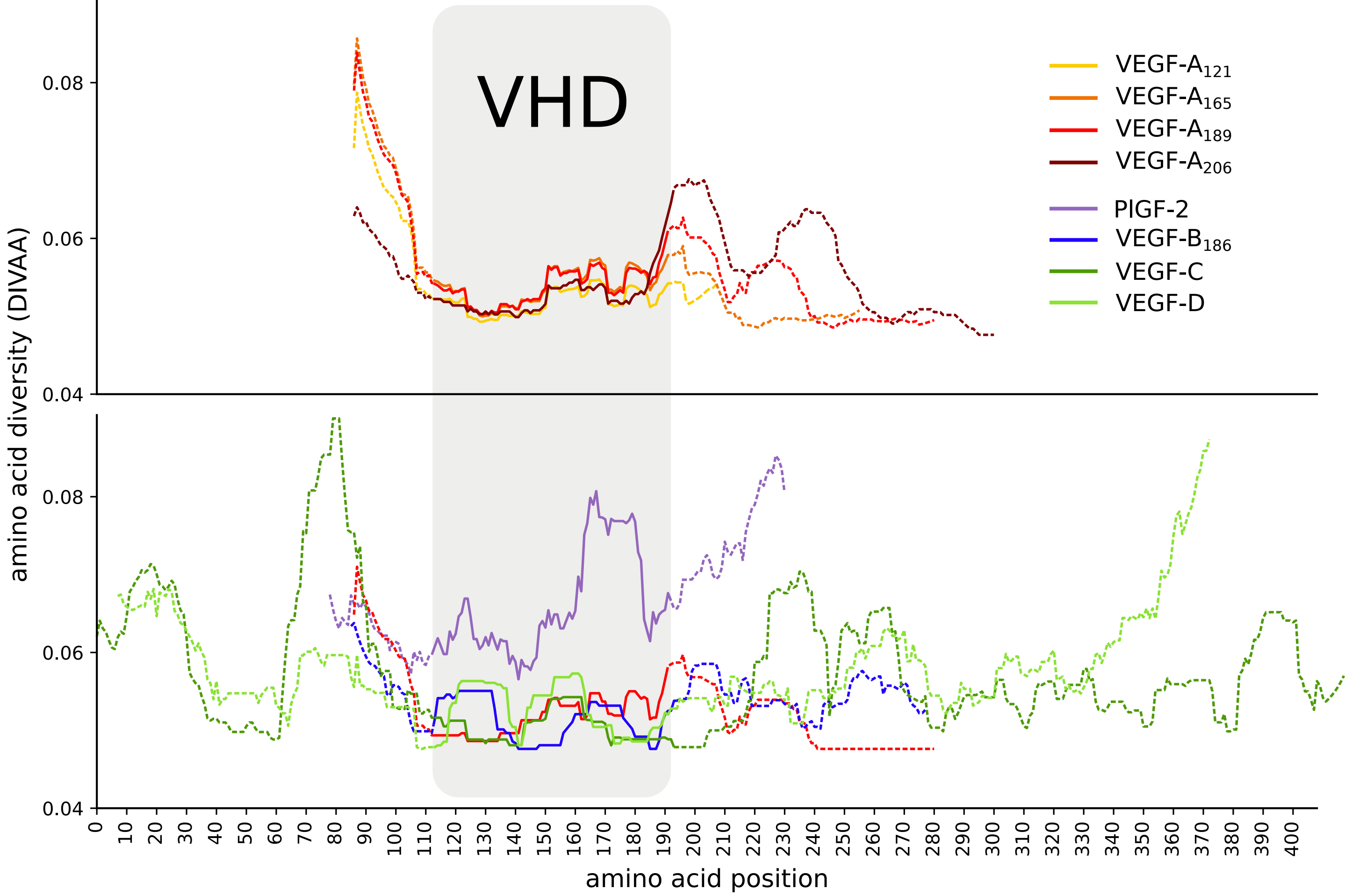

### Supplementary Figure 6

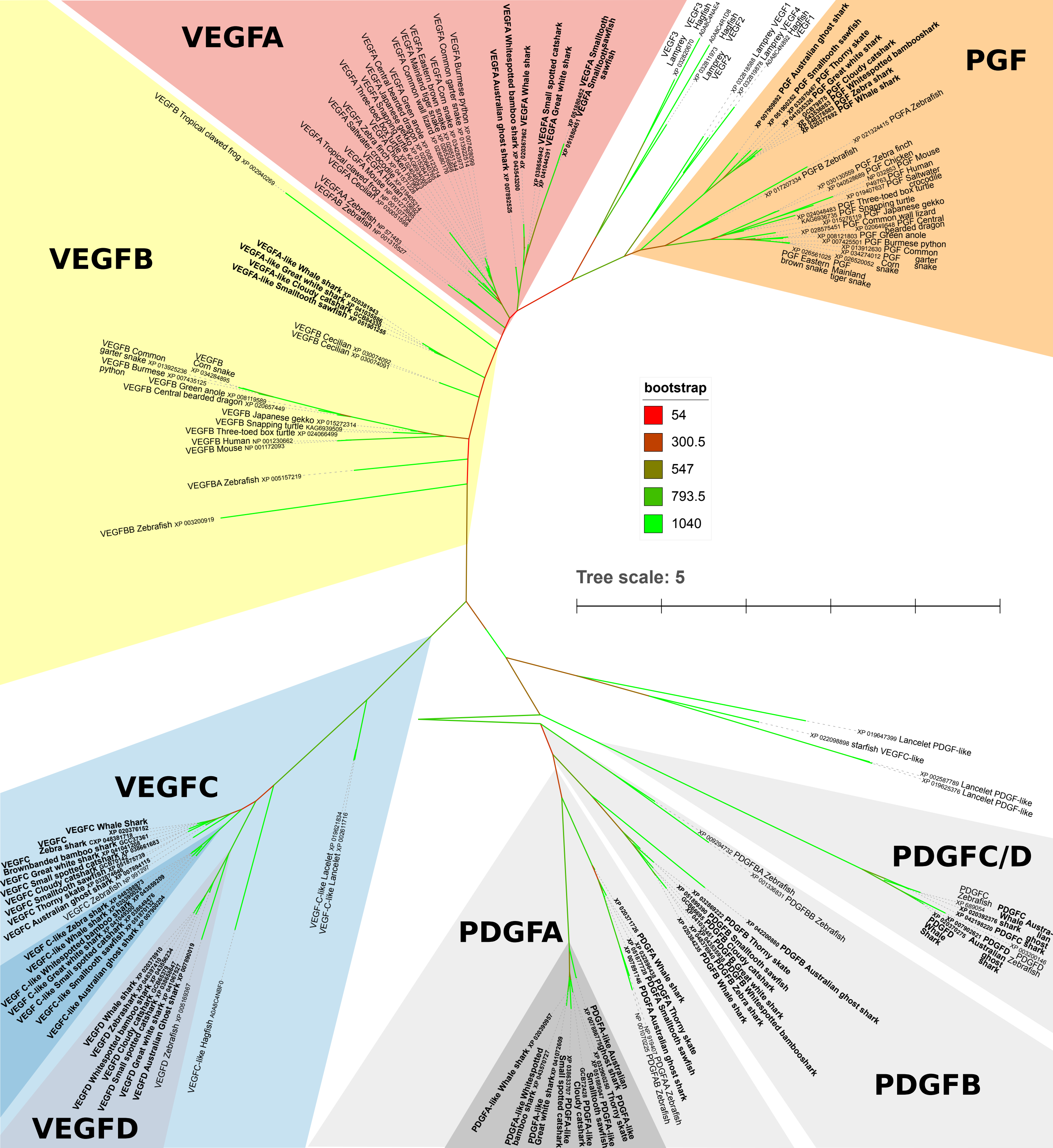

### Supplementary Figure 7

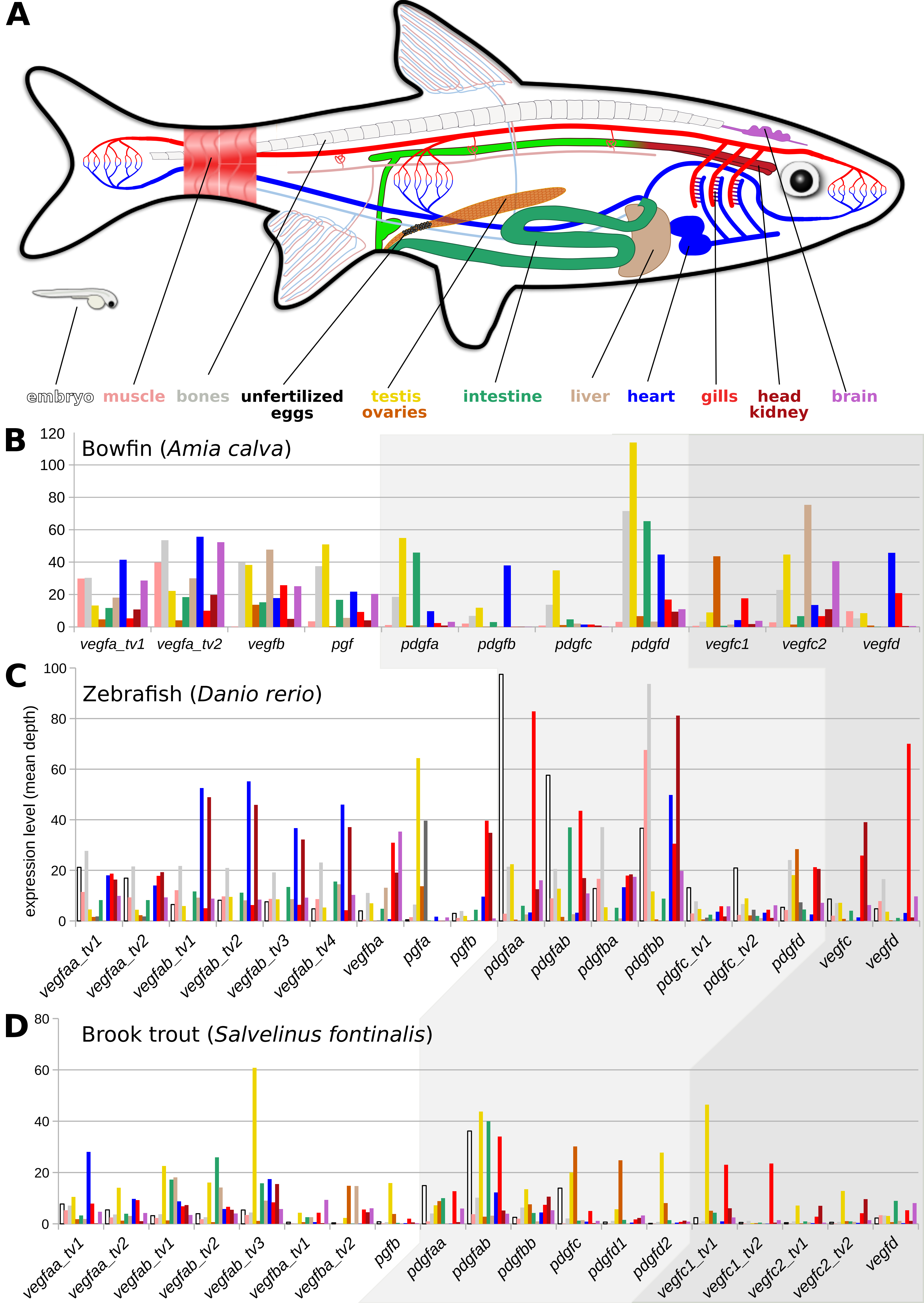
