## Supplementary Table 1 for "Expansion and collapse of VEGF diversity in major clades of the animal kingdom"

- Analysis performed: 190816_184756
- Analyzed sequences (hits resulting from 676 blast searches, 52 animal groups x 13 query sequences):5156 (out of which unique: 2379, programmatically recognized as VEGF/PDGF family members: 98.5%).
- Red dotted lines in the tree indicate paraphyletic relationships.
- The tree background color indicates the presence of the proteins with the corresponding color according to our hypotheses.
- The red-to-white background of the table indicates a heuristic reliability of the results, where a brighter color indicates a higher reliability. This is calculated using the number of fully sequenced genomes, the number of species in the phylum and the number of protein sequences available for that phylum.
- The numbers in the table denote the number of: orthologs found (black), P = paralogs found, ? = homologs found, whose relationship could not be programmatically determined, ∑ = total homologs found.

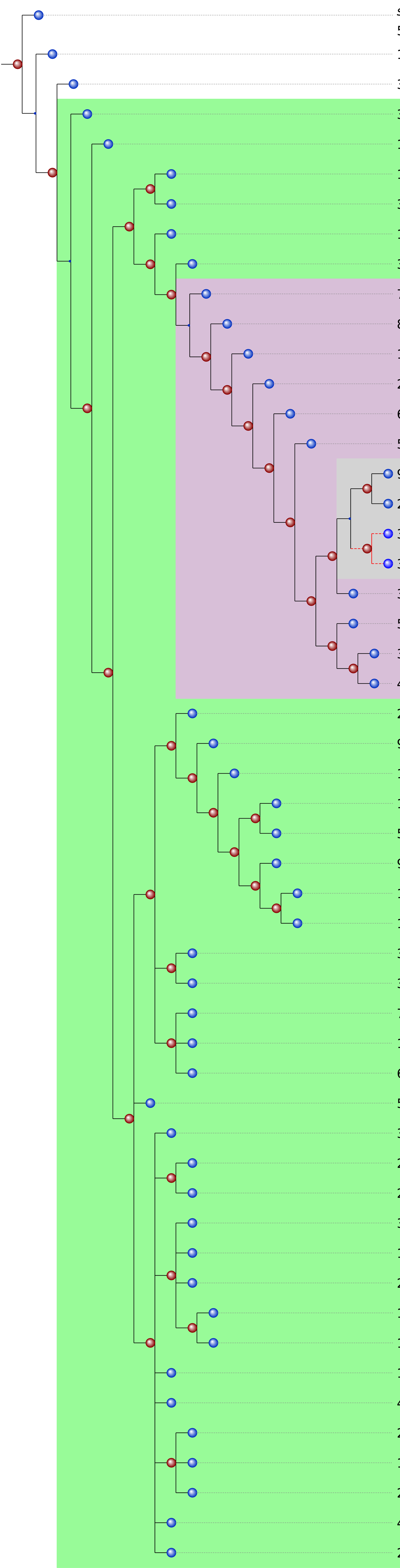

| # animal species | # sequences | # compl. genomes | # unique blasthits (excl. false pos.) | | PDGF-A | PDGF-B | PDGF-C | PDGF-D | PlGF-1 | VEGF-A121 | VEGF-A165 | VEGF-A206 | VEGF-B167 | VEGF-B186 | VEGF-C | VEGF-D | VEGF-F |
|---|---|---|---|---|---|---|---|---|---|---|---|---|---|---|---|---|---|
| 55 | 6 | 3 | 0 (0) | ctenophora (comb jellies) | | | | | | | | | | | | | |
| 1373 | 34k | 2 | 11 (0) | porifera (sponges) | | | | | | | | | | | 1 P0, ?7, ∑8 | 0 P1, ?1, ∑2 | 0 P0, ?3, ∑3 |
| 3 | 36k | 2 | 0 (0) | placozoa | | | | | | | | | | | | | |
| 3675 | 115k | 18 | 3 (3) | cnidaria (medusae/polyps) | 0 P6, ?4, ∑10 | 1 P3, ?1, ∑5 | 0 P1, ?0, ∑1 | | 0 P2, ?1, ∑3 | 0 P6, ?7, ∑13 | 1 P7, ?13, ∑21 | 0 P6, ?9, ∑15 | 0 P18, ?61, ∑79 | 0 P6, ?0, ∑6 | 11 P1, ?43, ∑55 | 0 P7, ?19, ∑26 | 0 P3, ?2, ∑5 |
| 151 | 925 | 0 | 1 (0) | xenacoelomorpha | | | | | | | | | | | | 0 P0, ?1, ∑1 | |
| 1797 | 136k | 11 | 3 (3) | echinodermata | 0 P5, ?1, ∑6 | 0 P5, ?0, ∑5 | 0 P11, ?1, ∑12 | 0 P12, ?1, ∑13 | 0 P9, ?3, ∑12 | 3 P2, ?7, ∑12 | 3 P3, ?11, ∑17 | 3 P2, ?7, ∑12 | 0 P10, ?9, ∑19 | 0 P10, ?8, ∑18 | 2 P8, ?12, ∑22 | 1 P9, ?7, ∑17 | 0 P7, ?3, ∑10 |
| 39 | 23k | 2 | 4 (4) | hemichordata (acorn wormws) | 0 P2, ?0, ∑2 | 0 P2, ?0, ∑2 | 0 P2, ?1, ∑3 | 0 P2, ?0, ∑2 | 0 P1, ?0, ∑1 | 0 P1, ?2, ∑3 | 0 P1, ?1, ∑2 | 0 P1, ?1, ∑2 | 0 P2, ?1, ∑3 | 0 P2, ?2, ∑4 | 2 P1, ?1, ∑4 | 0 P3, ?1, ∑4 | 0 P2, ?0, ∑2 |
| 11 | 95k | 4 | 7 (7) | cephalochordata (lancelets) | 0 P6, ?1, ∑7 | 0 P6, ?1, ∑7 | 0 P5, ?1, ∑6 | 0 P5, ?1, ∑6 | 0 P5, ?0, ∑5 | 1 P5, ?1, ∑7 | 1 P5, ?1, ∑7 | 1 P5, ?1, ∑7 | 0 P6, ?2, ∑8 | 0 P6, ?1, ∑7 | 6 P1, ?6, ∑13 | 0 P6, ?1, ∑7 | 0 P6, ?1, ∑7 |
| 362 | 64k | 6 | 2 (1) | tunicata | 0 P1, ?0, ∑1 | 0 P1, ?0, ∑1 | | | 0 P1, ?1, ∑2 | 1 P0, ?0, ∑1 | 1 P0, ?0, ∑1 | 1 P0, ?0, ∑1 | 0 P1, ?0, ∑1 | 0 P1, ?0, ∑1 | 0 P1, ?0, ∑1 | 0 P1, ?0, ∑1 | 0 P1, ?0, ∑1 |
| 77 | 8k | 3 | 0 (0) | cyclostomata (hagfish/lamprey) | | | | | | | | | | | | | |
| 833 | 115k | 6 | 52 (n.a.) | chondrichthyes (cartilaginous fishes) | 6 P21, ?0, ∑27 | 2 P25, ?0, ∑27 | 2 P18, ?0, ∑20 | 2 P11, ?0, ∑13 | 0 P25, ?0, ∑25 | 11 P14, ?0, ∑25 | 11 P14, ?0, ∑25 | 11 P13, ?0, ∑24 | 0 P29, ?0, ∑29 | 0 P29, ?1, ∑30 | 6 P23, ?0, ∑29 | 2 P26, ?0, ∑28 | 0 P25, ?0, ∑25 |
| 19097 | 2M | 186 | 674 (n.a.) | actinopterygii (ray-finned fishes) | 217 P665, ?170, ∑1052 | 27 P863, ?170, ∑1060 | 117 P282, ?15, ∑414 | 148 P269, ?9, ∑426 | 102 P882, ?159, ∑1143 | 426 P811, ?209, ∑1446 | 430 P462, ?137, ∑1029 | 433 P463, ?130, ∑1026 | 59 P1041, ?154, ∑1254 | 58 P1161, ?173, ∑1392 | 175 P1034, ?141, ∑1350 | 102 P1012, ?136, ∑1250 | 30 P1054, ?169, ∑1253 |
| 2 | 35k | 1 | 31 (30) | coelacanthimorpha (lobe-finned fishes) | 1 P15, ?0, ∑16 | 2 P12, ?0, ∑14 | 1 P12, ?0, ∑13 | 2 P4, ?0, ∑6 | 2 P12, ?0, ∑14 | 3 P12, ?0, ∑15 | 3 P11, ?0, ∑14 | 3 P11, ?0, ∑14 | 3 P14, ?0, ∑17 | 3 P14, ?1, ∑18 | 2 P15, ?0, ∑17 | 1 P16, ?0, ∑17 | 0 P14, ?0, ∑14 |
| 6 | 1k | 0 | 10 (8) | dipnoi (lungfishes) | 1 P3, ?0, ∑4 | 1 P3, ?0, ∑4 | 0 P4, ?0, ∑4 | 0 P4, ?0, ∑4 | 0 P4, ?0, ∑4 | 2 P2, ?0, ∑4 | 2 P2, ?0, ∑4 | | 0 P4, ?0, ∑4 | 0 P4, ?0, ∑4 | 0 P4, ?0, ∑4 | 0 P4, ?0, ∑4 | 0 P4, ?2, ∑6 |
| 5700 | 478k | 6 | 144 (n.a.) | amphibia | 16 P59, ?1, ∑76 | 12 P62, ?1, ∑75 | 6 P54, ?0, ∑60 | 5 P17, ?0, ∑22 | 0 P66, ?1, ∑67 | 27 P38, ?1, ∑66 | 27 P30, ?1, ∑58 | 27 P33, ?1, ∑61 | 3 P69, ?1, ∑73 | 3 P73, ?1, ∑77 | 5 P71, ?1, ∑77 | 6 P72, ?1, ∑79 | 0 P68, ?1, ∑69 |
| 9458 | 3M | 132 | 462 (n.a.) | aves (birds) | 127 P340, ?5, ∑472 | 118 P461, ?5, ∑584 | 98 P452, ?2, ∑552 | 154 P316, ?0, ∑470 | 109 P463, ?4, ∑576 | 140 P390, ?9, ∑539 | 141 P329, ?5, ∑475 | 141 P329, ?5, ∑475 | 0 P672, ?8, ∑680 | 0 P764, ?8, ∑772 | 130 P704, ?4, ∑838 | 96 P707, ?4, ∑807 | 1 P729, ?8, ∑738 |
| 24 | 179k | 4 | 102 (n.a.) | crocodylia (crocodiles) | 5 P26, ?0, ∑31 | 1 P14, ?0, ∑15 | 4 P35, ?0, ∑39 | 6 P21, ?0, ∑27 | 7 P46, ?0, ∑53 | 17 P40, ?0, ∑57 | 17 P35, ?0, ∑52 | 17 P29, ?0, ∑46 | 0 P57, ?0, ∑57 | 0 P63, ?0, ∑63 | 9 P54, ?0, ∑63 | 8 P55, ?0, ∑63 | 0 P53, ?0, ∑53 |
| 3249 | 96k | 6 | 65 (n.a.) | lepidosauria excl. toxicofera (non-poisonous lizards) | 3 P52, ?0, ∑55 | 6 P45, ?0, ∑51 | 3 P38, ?1, ∑42 | 5 P18, ?0, ∑23 | 6 P46, ?1, ∑53 | 26 P31, ?1, ∑58 | 26 P26, ?1, ∑53 | 26 P22, ?1, ∑49 | 2 P54, ?0, ∑56 | 2 P59, ?0, ∑61 | 7 P56, ?1, ∑64 | 5 P57, ?0, ∑62 | 4 P48, ?0, ∑52 |
| 3797 | 467k | 15 | 56 (n.a.) | toxicofera (poisonous reptiles) | 12 P122, ?0, ∑134 | 9 P113, ?0, ∑122 | 9 P89, ?0, ∑98 | 11 P26, ?0, ∑37 | 14 P91, ?28, ∑133 | 53 P63, ?29, ∑145 | 53 P43, ?29, ∑125 | 53 P45, ?28, ∑126 | 7 P109, ?26, ∑142 | 7 P120, ?26, ∑153 | 11 P108, ?26, ∑145 | 2 P119, ?23, ∑144 | 24 P107, ?5, ∑136 |
| 358 | 184k | 10 | 70 (n.a.) | testudines (turtles) | 9 P65, ?1, ∑75 | 8 P66, ?1, ∑75 | 7 P60, ?7, ∑74 | 17 P20, ?0, ∑37 | 10 P61, ?1, ∑72 | 30 P48, ?1, ∑79 | 30 P52, ?4, ∑86 | 30 P36, ?1, ∑67 | 8 P65, ?1, ∑74 | 8 P88, ?2, ∑98 | 6 P91, ?2, ∑99 | 5 P92, ?1, ∑98 | 0 P70, ?1, ∑71 |
| 5 | 26k | 1 | 25 (25) | monotremata (egg-laying mammals) | 1 P6, ?0, ∑7 | 1 P5, ?0, ∑6 | 3 P10, ?0, ∑13 | 3 P10, ?0, ∑13 | 1 P5, ?0, ∑6 | 1 P6, ?0, ∑7 | 1 P6, ?0, ∑7 | 1 P6, ?0, ∑7 | 0 P8, ?0, ∑8 | 0 P8, ?0, ∑8 | 2 P6, ?0, ∑8 | 1 P7, ?0, ∑8 | 0 P6, ?0, ∑6 |
| 334 | 142k | 5 | 88 (n.a.) | metatheria (marsupials) | 7 P36, ?0, ∑43 | 5 P36, ?0, ∑41 | 4 P33, ?0, ∑37 | 10 P16, ?0, ∑26 | 4 P25, ?0, ∑29 | 4 P28, ?0, ∑32 | 4 P22, ?0, ∑26 | 4 P22, ?0, ∑26 | 4 P32, ?0, ∑36 | 4 P36, ?0, ∑40 | 4 P35, ?0, ∑39 | 5 P34, ?0, ∑39 | 0 P39, ?0, ∑39 |
| 4781 | 8M | 181 | 509 (n.a.) | eutheria (placentals) | 247 P762, ?6, ∑1015 | 223 P892, ?13, ∑1128 | 218 P659, ?1, ∑878 | 235 P417, ?0, ∑652 | 261 P1262, ?9, ∑1532 | 434 P900, ?6, ∑1340 | 440 P862, ?6, ∑1308 | 440 P857, ?6, ∑1303 | 249 P1420, ?11, ∑1680 | 249 P1504, ?10, ∑1763 | 171 P1406, ?9, ∑1586 | 164 P1601, ?9, ∑1774 | 0 P1596, ?10, ∑1606 |
| 200 | 46k | 2 | 3 (3) | tardigrada (water bears) | 0 P1, ?3, ∑4 | 0 P3, ?1, ∑4 | 0 P2, ?0, ∑2 | 0 P1, ?0, ∑1 | 0 P2, ?0, ∑2 | 0 P0, ?4, ∑4 | 0 P0, ?6, ∑6 | 0 P0, ?3, ∑3 | 0 P4, ?0, ∑4 | 0 P3, ?1, ∑4 | 0 P2, ?2, ∑4 | 0 P4, ?0, ∑4 | 0 P2, ?1, ∑3 |
| 96 | 2k | 1 | 0 (0) | onychophora (velvet worms) | | | | | | | | | | | | | |
| 195 | 2k | 0 | 0 (0) | pycnogonida (sea spiders) | | | | | | | | | | | | | |
| 10263 | 646k | 27 | 11 (11) | arachnida (spiders) | 2 P20, ?3, ∑25 | 0 P20, ?3, ∑23 | 0 P21, ?5, ∑26 | 0 P9, ?2, ∑11 | 2 P24, ?39, ∑65 | 14 P5, ?22, ∑41 | 14 P5, ?20, ∑39 | 13 P5, ?16, ∑34 | 0 P25, ?13, ∑38 | 0 P24, ?11, ∑35 | 1 P16, ?16, ∑33 | 1 P14, ?2, ∑17 | 1 P26, ?30, ∑57 |
| 5 | 39k | 1 | 11 (10) | xiphosura (horseshoe crabs) | 0 P6, ?3, ∑9 | 0 P6, ?1, ∑7 | 0 P7, ?3, ∑10 | 0 P3, ?2, ∑5 | 0 P7, ?20, ∑27 | 7 P0, ?11, ∑18 | 7 P0, ?11, ∑18 | 7 P0, ?11, ∑18 | 0 P7, ?11, ∑18 | 0 P7, ?11, ∑18 | 0 P7, ?4, ∑11 | 0 P2, ?0, ∑2 | 0 P7, ?11, ∑18 |
| 978 | 7k | 1 | 1 (0) | myriapoda (millipeds) | | | | | | | | | | 0 P0, ?1, ∑1 | | | |
| 10996 | 947k | 25 | 3 (3) | crustacea | 0 P10, ?8, ∑18 | 0 P11, ?4, ∑15 | 1 P4, ?3, ∑8 | 0 P1, ?8, ∑9 | 0 P5, ?7, ∑12 | 5 P4, ?11, ∑20 | 5 P4, ?12, ∑21 | 4 P4, ?8, ∑16 | 1 P9, ?19, ∑29 | 1 P6, ?19, ∑26 | 1 P8, ?13, ∑22 | 1 P6, ?8, ∑15 | 1 P7, ?9, ∑17 |
| 114720 | 7M | 339 | 2 (2) | hexapoda (insects) | 20 P77, ?117, ∑214 | 3 P89, ?113, ∑205 | 1 P42, ?34, ∑77 | 2 P23, ?16, ∑41 | 3 P52, ?141, ∑196 | 49 P53, ?154, ∑256 | 59 P61, ?198, ∑318 | 39 P45, ?129, ∑213 | 5 P110, ?177, ∑292 | 5 P94, ?156, ∑255 | 16 P18, ?98, ∑132 | 2 P36, ?55, ∑93 | 3 P95, ?236, ∑334 |
| 30 | 368 | 0 | 0 (0) | nematomorpha (horsehair worms) | | | | | | | | | | | | | |
| 3524 | 2M | 100 | 1 (1) | nematoda (roundworms) | 0 P0, ?35, ∑35 | 0 P0, ?11, ∑11 | | | | | | | 0 P1, ?22, ∑23 | 0 P0, ?21, ∑21 | 0 P0, ?4, ∑4 | 0 P0, ?2, ∑2 | 0 P0, ?24, ∑24 |
| 7 | 21k | 1 | 8 (1) | priapulida (penis worms) | 1 P0, ?0, ∑1 | 0 P1, ?7, ∑8 | | | | | 0 P1, ?0, ∑1 | | 0 P1, ?0, ∑1 | 0 P1, ?0, ∑1 | 0 P1, ?0, ∑1 | | 0 P1, ?0, ∑1 |
| 1 | 1 | 0 | 0 (0) | loricifera | | | | | | | | | | | | | |
| 62 | 436 | 0 | 0 (0) | kinorhyncha (mud dragons) | | | | | | | | | | | | | |
| 56 | 2k | 0 | 0 (0) | chaetognatha (arrow worms) | | | | | | | | | | | | | |
| 327 | 3k | 0 | 1 (0) | bryozoa (moss animals) | | | | | | | | | | 0 P0, ?1, ∑1 | | | |
| 26 | 155 | 0 | 2 (0) | entoprocta | | | | | | | | | | 0 P0, ?1, ∑1 | | 0 P0, ?1, ∑1 | |
| 2 | 278 | 0 | 0 (0) | cycliophora (symbion) | | | | | | | | | | | | | |
| 3338 | 129k | 5 | 1 (1) | annelida (segmented worms) | 0 P0, ?3, ∑3 | 0 P2, ?2, ∑4 | 0 P1, ?1, ∑2 | 0 P1, ?1, ∑2 | 0 P1, ?2, ∑3 | 0 P0, ?3, ∑3 | 0 P0, ?3, ∑3 | 0 P0, ?3, ∑3 | 0 P1, ?2, ∑3 | 0 P1, ?1, ∑2 | 0 P0, ?3, ∑3 | 0 P1, ?1, ∑2 | 0 P1, ?2, ∑3 |
| 14215 | 742k | 26 | 2 (2) | mollusca | 0 P4, ?6, ∑10 | 0 P8, ?0, ∑8 | 0 P4, ?4, ∑8 | 0 P1, ?4, ∑5 | 0 P2, ?1, ∑3 | 2 P2, ?9, ∑13 | 2 P2, ?9, ∑13 | 2 P2, ?5, ∑9 | 0 P9, ?6, ∑15 | 0 P4, ?6, ∑10 | 0 P1, ?15, ∑16 | 0 P7, ?4, ∑11 | 0 P8, ?4, ∑12 |
| 262 | 5k | 1 | 0 (0) | nemertea (ribbon worms) | | | | | | | | | | | | | |
| 100 | 42k | 1 | 6 (2) | brachiopoda (lamp shells) | 0 P1, ?0, ∑1 | 0 P1, ?0, ∑1 | 0 P0, ?1, ∑1 | 0 P1, ?0, ∑1 | | 0 P1, ?0, ∑1 | 0 P1, ?0, ∑1 | 0 P1, ?0, ∑1 | 0 P1, ?1, ∑2 | 0 P1, ?0, ∑1 | 0 P1, ?2, ∑3 | 1 P0, ?0, ∑1 | 0 P1, ?0, ∑1 |
| 14 | 165 | 1 | 0 (0) | phoroniformea (horseshoe worms) | | | | | | | | | | | | | |
| 130 | 389 | 0 | 0 (0) | gastrotricha (hairybacks) | | | | | | | | | | | | | |
| 4472 | 561k | 26 | 1 (0) | platyhelminthes (flatworms) | | | | | | | | | | 0 P0, ?1, ∑1 | | | |
| 21 | 79 | 0 | 0 (0) | gnathostomulida (jaw worms) | | | | | | | | | | | | | |
| 1 | 2 | 0 | 0 (0) | micrognathozoa | | | | | | | | | | | | | |
| 237 | 64k | 6 | 6 (2) | rotifera (wheel animals) | 1 P0, ?0, ∑1 | 0 P1, ?0, ∑1 | 0 P1, ?1, ∑2 | | 0 P1, ?0, ∑1 | | 0 P1, ?0, ∑1 | | 0 P1, ?0, ∑1 | 0 P1, ?1, ∑2 | 0 P1, ?0, ∑1 | 0 P1, ?0, ∑1 | 0 P1, ?2, ∑3 |
| 4 | 9k | 1 | 1 (0) | orthonectida | | | | | | | | | | | 0 P0, ?1, ∑1 | | |
| 24 | 150 | 0 | 0 (0) | dicyemida | | | | | | | | | | | | | |

Force topology is enabled!
Branch lengths do not represent real values.
